# Direction of actin flow dictates integrin LFA-1 orientation during leukocyte migration

**DOI:** 10.1101/071936

**Authors:** Pontus Nordenfelt, Travis I. Moore, Shalin B. Mehta, Joseph Mathew Kalappurakkal, Vinay Swaminathan, Nobuyasu Koga, Talley J. Lambert, David Baker, Jennifer C. Waters, Rudolf Oldenbourg, Tomomi Tani, Satyajit Mayor, Clare M. Waterman, Timothy A. Springer

## Abstract

Integrin αβ heterodimer cell surface receptors mediate adhesive interactions that provide traction for cell migration. Here, we test whether the integrin head, known from crystal structures, adopts a specific orientation dictated by the direction of actin flow on the surface of migrating cells. We insert GFP into the rigid head of the full integrin, model with Rosetta the orientation of GFP and its transition dipole relative to the integrin, and measure orientation with fluorescence polarization microscopy. Dependent on coupling to the cytoskeleton, integrins orient in the same direction as retrograde actin flow with their cytoskeleton-binding β-subunits tilted by applied force. The measurements demonstrate that intracellular forces can orient cell surface integrins and support a molecular model of integrin activation by cytoskeletal force. We have developed a method that places atomic, ~Å structures of cell surface receptors in the context of functional, cellular length-scale, ~μm measurements and shows that rotation and tilt of cell surface receptors relative to the membrane plane can be restrained by interactions with other cellular components.

## Introduction

The integrin lymphocyte function-associated antigen-1 (LFA-1, αLβ2) participates in a wide range of adhesive interactions including antigen recognition, emigration from the vasculature, and migration of leukocytes within tissues^1,2^. Integrin ectodomains assume three global conformational states (Fig. 1a) with the extended-open conformation binding ligand with ~1,000-fold higher affinity than the bent-closed and extended-closed conformations^2^. Binding of LFA-1 to intercellular adhesion molecule (ICAM) ligands by the αI domain in the integrin head is communicated through the β-subunit leg, transmembrane, and cytoplasmic domains to the actin cytoskeleton via adaptors such as talins and kindlins that bind specific sites in the β-subunit cytoplasmic domain^3^. As reviewed^4,5^, measurements of traction force on substrates and more specific measurements of force within ligands and cytoskeletal components have suggested that integrins are important in force transmission between extracellular ligands and the actin cytoskeleton. Forces on the cytoplasmic domain of the LFA-1 β_2_-subunit have been measured in the 1-6 pN range and associated with binding to ligand and the cytoskeleton^6^.

**Figure 1.**
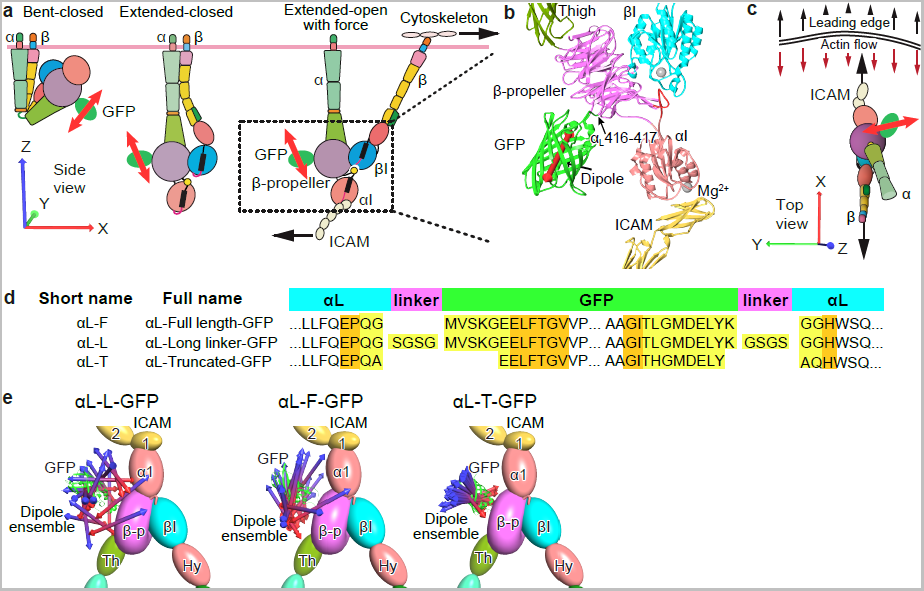
GFP-LFA-1 fusions, emission anisotropy, and integrin head segmental motion. **a.** Three global conformational states of integrins^2^. Cartoons depict each integrin domain and GFP with its transition dipole (red double-headed arrows). **b.** Ribbon diagram of the integrin headpiece of αL-T bound to ICAM-1. The GFP insertion site in the β-propeller domain is arrowed. Dipole is shown in red. **c.** Cartoon as in (a) of ICAM-engaged, extended-open LFA-1 showing direction of leading edge motion and actin flow. Large arrows show pull on integrin-β by actin and resistance by ICAM-1. Axes shown in (a) and (c) are similar to those in the references state in Fig. 6. **d.** Sequences and boundaries used in GFP-LFA-1 fusions. Highlighted residues were modeled by Rosetta to link GFP to the integrin (yellow) and altered in backbone and sidechain orientation to minimize energy (yellow and orange). **e.** Orientation of the transition dipole in GFP-LFA-1 fusions. Integrin domains are shown as ellipsoids or torus and GFP is shown in cartoon for 1 ensemble member. GFP transition dipoles are shown as cylinders with cones at each end for 20 representative Rosetta ensemble members, with the asymmetry of GFP referenced by using different colors for the ends of transition dipoles (which themselves have dyad symmetry).

Tensile forces exerted through integrins between the actin cytoskeleton and extracellular ligands have the potential to align integrins; such alignment would in turn provide the opportunity to test alternative models of integrin activation. Some models suggest that binding of the cytoskeletal adaptor protein talin to the integrin β-subunit cytoplasmic domain is fully sufficient to activate high affinity of the extracellular domain for ligand^7,8^. Other models, supported by steered molecular dynamics and measurements in migrating cells, have proposed that tensile force stabilizes the high-affinity, extended-open integrin conformation because of its increased length along the tensile force-bearing direction compared to the other two integrin conformations (Fig. 1a)^6,9,,-12^. These models propose that inherent in the three conformational states of integrins is a mechanism by which integrin adhesiveness can be activated when the integrin simultaneously binds the actin cytoskeleton and an extracellular ligand that can resist cytoskeleton-applied force. Thus, the same intracellular effectors that regulate actin dynamics can simultaneously and coordinately regulate cell adhesion to provide the traction for cellular chemotaxis and migration. Furthermore, directional migration is a critical aspect of immune cell function, and alignment of integrins by activation would provide a mechanism for directional sensing. While this model is appealing to structural biologists, it requires validation in cells.

Here, we test a key prediction of the cytoskeletal force model of integrin activation: that the tensile force exerted through integrins between the actin cytoskeleton and extracellular ligands as they function in cell migration causes them to assume a specific orientation and tilt on the cell surface relative to the direction of pulling on the integrin by actin retrograde flow; actin flow is known to be locally aligned for migrating fibroblast and epithelial cells^13,14^. Measuring integrin orientation on cell surfaces also provides an opportunity to correlate crystal structures of integrins at the Å length scale with microscopic measurements on integrin-bearing cells at the micron length scale. Integrating measurements at such different length scales is a long-standing goal of many fields of biological research. While integrins like other membrane proteins are generally free to rotate in the plane of the membrane, tensile force would cause an integrin to orient in the same direction as the pulling force. Like most membrane proteins, integrins are drawn in cartoons (as in Fig. 1a) as projecting with their leg-like domains normal to the plasma membrane; however, resting integrins are free to tilt^15^ and force could tilt the integrin far from the membrane normal. In general, despite a wealth of structures for membrane protein ectodomains, little is known about their orientation on cell surfaces.

In this work, we make use of previous structural studies on integrins^2,16,17^, and orient these structures in a reference frame that corresponds to the plasma membrane of a migrating lymphocyte. In addition to general structural knowledge on many integrin families, we make specific use of crystal structures for the αI domain of LFA-1 bound to ICAMs, the LFA-1 headpiece, and two states of the bent ectodomain of the LFA-1 (αLβ2) relative, αXβ2. We also use negative stain EM class averages showing the bent-closed, extended-closed, and extended-open conformations of the αLβ2 and αXβ2 ectodomains. These structures together with those of green fluorescent protein (GFP) have guided development here of constrained integrin-GFP fusions and prediction using Rosetta^18^ of the orientation of GFP and its fluorescent excitation/emission transition dipole relative to the integrin. Two different types of fluorescent microscopes provide similar measurements of the orientation of the transition dipole relative to the direction of actin flow. Integrin–ligand engagement in combination with cytoskeletal force results in spatially ordered organization of LFA-1 in the protrusive lamellipodial region and is dependent on the movement vector of the underlying actin cytoskeletal framework^19,20^. The results show that actin flow from the leading edge dictates a specific molecular orientation on the cell surface of LFA-1 and support the cytoskeletal force model of integrin activation.

## Results

### Design, simulation, and testing of constrained LFA-1- GFP

To report integrin orientation on cell surfaces, we inserted GFP into a loop of the integrin β-propeller domain (Fig. 1a-d). This allows monitoring of the orientation of both the β-propeller and βI domains, which come together over a large, highly stable, rigid interface to form the integrin head. The β-propeller was chosen because of its rigid structure, its lack of participation in integrin conformational change^2^, and the availability of a previously validated insertion position that is remote from other integrin domains in all three conformational states^21^. We tested multiple fusions, including one in which residues were added to increase flexibility, and those that deleted residues from N and C-terminal segments of GFP that are disordered or vary in position among GFP crystal structures and were designed to constrain GFP orientation (Supplemental Table 1). We modeled with Rosetta any introduced linker residues, residues that vary in position in independent GFP structures, and residues in LFA-1 adjacent to the inserted GFP (Fig. 1d). A wide range of possible orientations of the two connections between LFA-1 and GFP was effectively sampled using polypeptide segments from the protein databank, selecting those that enabled connections at both the N and C-termini of GFP to be closed, and then minimizing the energy of the system with respect to the degrees of freedom of the connecting linkers^18^. The distribution of dipole orientations in the resulting ensembles (Fig. 1e) provides a range of orientations in which the actual orientation should be included, and may approximate the contribution of variation in GFP-integrin orientation to a decrease in emission anisotropy measurable with fluorescence polarization microscopy.

We used emission anisotropy total internal reflection fluorescence microscopy (EA-TIRFM)^22^ (Fig. 2a) to examine GFP-LFA-1 fusions stably transfected in Jurkat T lymphocytes that were allowed to randomly migrate on ICAM-1 substrates in the xy focal plane of the microscope (Fig. 2b). In EA-TIRFM, s-polarized light excites fluorophores that are parallel to the plane of incoming light, and the parallel and perpendicular components of the emission fluorescence are recorded on separate cameras (Fig. 2a). The excitation and emission dipoles of GFP are highly aligned and may be referred to collectively as the transition dipole^23,24^. Furthermore, the difference in time between excitation and emission is short relative to tumbling time for GFP ^22^ (and integrin-GFP fusions). Our measurements of fluorescence intensity are for the ensemble of LFA-1-GFP fusions within individual pixels (Fig. 2b); thus for constrained GFP-LFA-1 fusions that are similarly oriented (Fig. 2c), more emitted fluorescence will be recorded on the parallel than perpendicular camera, resulting in high emission anisotropy (Fig. 2b). In contrast, variation in GFP orientation relative to the integrin in unconstrained GFP (Fig. 2d) or random orientation GFP-LFA-1 fusions in the plane of the membrane (Fig. 2b, pixel 1) will lead to a relative increase in the intensity recorded in the perpendicular camera, resulting in lower emission anisotropy (Fig. 2a).

**Figure 2.**
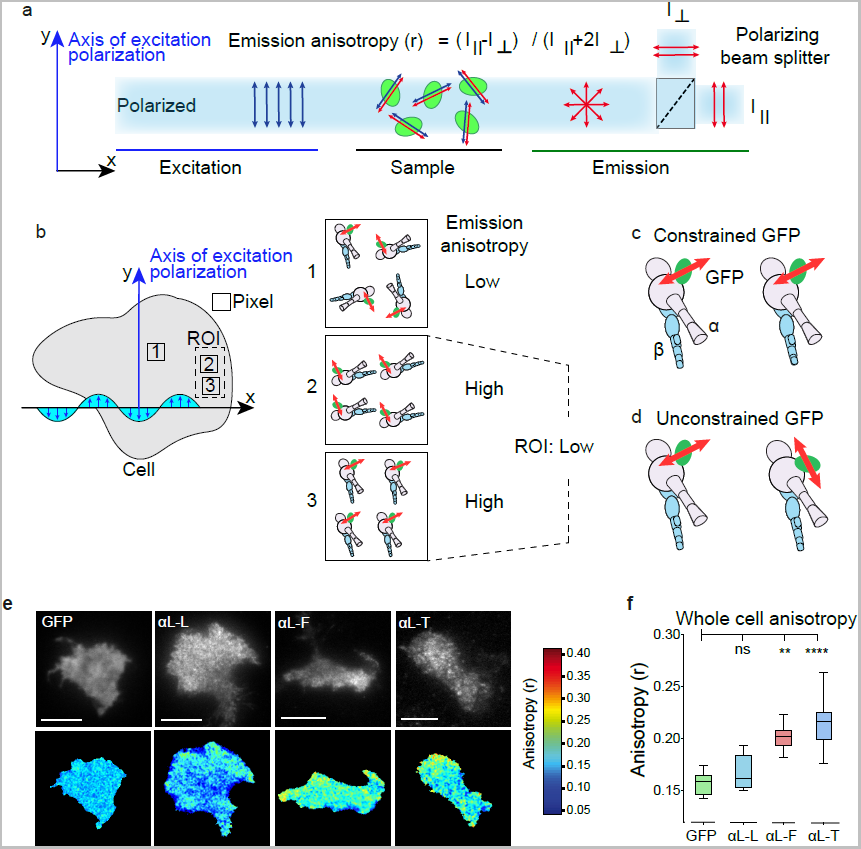
Emission anisotropy of GFP-LFA-1 fusions. **a.** Schematic of Emission Anisotropy TIRF Microscopy. The transmission dipole of GFP has an excitation dipole (blue) that is very close in orientation to its emission dipole (red). **b.** Schematic showing a cell in same microscope xy plane as in (a) and different outcomes in emission anisotropy when using a constrained integrin-GFP fusion. Depending on the orientation of integrins within pixels (1-3) or across pixels (region of interest, ROI), the emission anisotropy will change as indicated. **c-d.** Schematic of aligned integrins with constrained (c) or unconstrained GFP (d). **e.** Representative images from movies of Jurkat T cells stably expressing GFP-LFA-1 fusions migrating on ICAM-1. Each pair of panels shows total GFP fluorescence intensity (upper) and anisotropy (lower, color scale bar to right). Scale bars are 5 μm. **f.** Emission anisotropy of GFP-LFA-1 fusions, averaged over Jurkat T cells migrating in random directions, in at least five independent experiments. Box plots show the full range (whiskers) of observations with median as line and 25-75 percentile range boxed. Kruskal-Wallis test with multiple comparison correction gave the indicated p-values. N (number of cells) from left was 18, 22, 17, 37. *: p < 0.05; **: p < 0.01; ****: p < 0.0001.

To test the Rosetta modeling predictions of GFP constrainment, three GFP integrin fusions were studied on the surface of live Jurkat T lymphoblasts migrating on ICAM-1 substrates (Fig. 2e). Emission anisotropy was highest for the truncated construct, αL-T; intermediate for the full length construct, αL-F; and lowest for the construct with added linker residues, αL-L, which had an emission anisotropy comparable to cytoplasmic GFP (Fig. 2e-f). Measured anisotropy was consistent with the predicted GFP transition dipole alignment among members of structural ensembles obtained via Rosetta (Fig. 1e). Anisotropies of aL-F and aL-T-GFP that are substantially higher than for GFP imply that the integrins are not oriented randomly about an axis normal to the plane of membrane, but are aligned with one another in individual pixels (Fig. 2b, pixels 2 and 3); however, integrin orientation might nonetheless differ between pixels (Fig. 2b, ROI). The lack of alignment seen with αL-L compared to the increasing alignment seen with αL-F and αL-T suggests that the orientation of the GFP transition dipole in αL-L is not correlated with the orientation of the integrin (Fig. 2d) as seen in αL-L Rosetta ensembles (Fig. 1e). At high fluorophore concentrations, anisotropy can be reduced by homo-FRET^22^. However, LFA-1-GFP anisotropy was independent of fluorescence intensity, showing little or no homo-FRET (Supplemental Fig. 1).

### Quantitative description of actin flow in migrating T-cells

When integrins are bound to ligand, we predict that traction forces act on the integrin β-subunit cytoplasmic domain and dictate local integrin alignment. A strong candidate for this force is actin retrograde flow generated through actin filament extension along the membrane at the cell front ^25^. Before determining the orientation of our LFA-1-GFP fusions relative to actin flow, it was necessary to measure actin flow in migrating lymphocytes. Actin flow is known to be retrograde and normal to the leading edge in migrating fibroblasts and epithelial cells^13,14,26^. Much less is known about actin flow in migrating lymphocytes, although it is known to be retrograde within the immunological synapse in non-migrating T cells^27,28^. We defined actin dynamics in migrating T cells expressing lifeact-mNeonGreen using the super-resolution capabilities of structured illumination microscopy (SIM). Actin flow velocity and direction were determined using optical flow analysis^29,30^ of the texture maps generated by time lapse SIM along the leading edge relative to the membrane. Migrating T cells lacked organized actin stress fibers characteristic of many other cells, such as epithelial cells and fibroblasts (Fig. 3a). Optical flow analysis of time-lapse movies (Fig. 3a) showed that flow was retrograde at close to 90° relative to the leading edge (Fig. 3b) with a velocity of 17 ± 1 nm/s on ICAM-1 substrates (Fig. 3c, Movie 3) as confirmed and consistent with kymographic analysis^6,31^ (Supplemental Fig. 2). On the non-integrin substrate, anti-CD43, actin flow was significantly faster (71 ± 4 nm/sec, Movie 4) compared to ICAM-1, while flow on mixed ICAM-1 and anti-CD43 substrates was intermediate in velocity (34 ± 2 nm/sec) (Fig. 3c, Movie 5).

**Figure 3.**
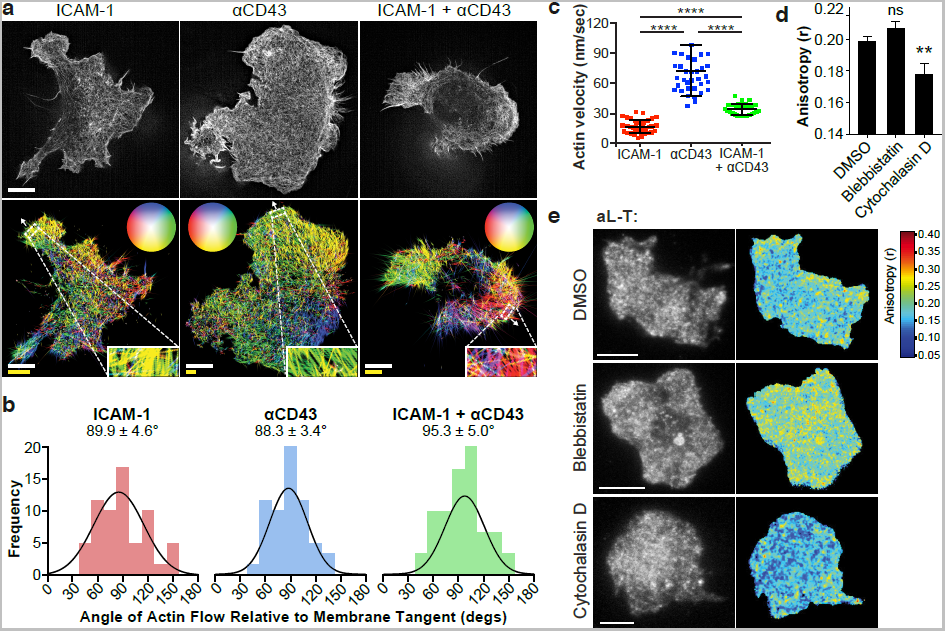
Actin flow dynamics, orientation, and relation to GFP-LFA-1 anisotropy in migrating T cells. **a.** Representative frames of structured illumination microscopy movies (above) of the actin cytoskeleton visualized with lifeact-mNeonGreen in Jurkat T cells migrating on ICAM-1 (10 μg/ml), anti-CD43 IgG (10 μg/ml), or their mixture (10 μg/ml each). Optical actin flow vector maps of the same cells (below) with zoom insets of representative areas analyzed. Vectors encode flow direction by color (circular keys in the direction from center of circle to perimeter) and velocity by length. Scale bars encode dimensions in the micrograph (white, 5 μm) and velocity (yellow, 30 nm/s). **b.** Actin flow direction relative to tangent of leading edge membrane. Bins of 15° are shown with Gaussian fit and mean ± SEM. ICAM-1, N=62; αCD43, N=63; ICAM-1+αCD43, N=67. **c.** Leading edge actin flow velocity from optical flow analysis. Plots show full range of the data. Bars show mean ± SD. Two-tailed Mann-Whitney tests all show p < 0.0001 (****). ICAM-1, N=39; αCD43 N=38; and ICAM-1+ αCD43, N=25. **d, e.** Jurkat T cells migrating on ICAM-1 (10 μg/ml) expressing αL-T-GFP were treated with DMSO as control, blebbistatin (100 μM), or cytochalasin D (100 nM) and fixed. d. Whole cell emission anisotropy analyzed as in Fig. 2c. Mann-Whitney test for DMSO (N=37) vs blebbistatin (N=9) was 0.130 and for DMSO vs cytochalasin D (N=14) was 0.003. ** p < 0.01. **e.** Representative total fluorescence intensity (I|| + 2I⊥, top) and anisotropy (bottom). Scale bars: 5 μm.

The ability of ICAM-1 on substrates to slow retrograde actin flow, and the intermediate results with mixed anti-CD43 and ICAM-1 substrates, strongly suggest that LFA-1 is mechanically linked to the actin in retrograde flow. Activating LFA-1 independently of intracellular signaling with Mn^2+^ treatment increased actin flow velocity (28 ± 2 nm/sec) relative to untreated cells (Supplemental Fig. 2), suggesting that artificial activation with Mn^2+^ disrupts integrin association with actin. Linkage observed here between LFA-1 and the actin cytoskeleton to ICAM-1 immobilized on a glass coverslip is in agreement with previous studies on immunological synapses formed on bilayers in which ICAM-1 can diffuse and be dragged by LFA-1^27,28^. However, actin flow in lymphocytes on lipid bilayers is more rapid at ~100- 300 nm/s, and the ICAM-1 is dragged along at about 40% of this rate, demonstrating a clutch-like connection. Although indirect regulatory mechanisms cannot be ruled out, the most straightforward interpretation of our results is that flowing actin is slowed by the linkage of LFA-1 to both the actin cytoskeleton inside the cell and ICAM-1 on the substrate. Linkage of engaged LFA-1 to a force-producing actin network is similar to that proposed in the molecular clutch model of mesenchymal cell migration^32,33^.

We further tested the role of the actin cytoskeleton in integrin alignment by disrupting two prominent drivers of actin flow, contractility and polymerization. Blebbistatin, an inhibitor of myosin-dependent actin filament contractility, had no effect on αL-T anisotropy (Fig. 3d-e), consistent with the lack in T cells of actin stress fibers. In contrast, cytochalasin D, an inhibitor of actin polymerization, significantly decreased LFA-1 anisotropy (Fig. 3d-e). Together, these results show that an intact cytoskeleton is required for LFA-1 anisotropy and suggest that actin polymerization, a mechanism operative in the lamellipodium to generate retrograde flow that is independent of actomyosin contraction^13^, is important for LFA-1 anisotropy and hence alignment.

### Two independent polarization microscopy techniques give similar LFA-1 orientations near the migratory leading edge

Having shown that αL-T and αL-F LFA-1-GFP fusions were aligned within individual pixels, we next determined whether integrins in larger regions near cell leading edges were aligned with one another, and whether this alignment correlated with the direction of actin flow, a measure that we term “orientation.” To do this we employed two independent microscopy techniques.

We first used the photoselective properties of EA-TIRFM^22^ to measure orientation. Fluorescence anisotropy shows cos^2^ dependence on the angle between the electric field of the polarized light and the fluorophore transition dipole^34,35^, where A represents the amplitude in angular dependence of anisotropy, γ the angle between the membrane normal and excitation axis, and θ_d_ the angle between the transition dipole and the membrane normal (Fig. 4a, Methods Eq. 5). We verified this dependence by imaging actin filaments, either assembled in vitro and stained with SiR-actin or in migrating T cells stained with Alexa488-phalloidin. We defined γ and θ_d_ relative to the long axis of actin filaments instead of the membrane normal. Fits of the cos^2^ dependence of anisotropy gave θ_d_ = 86° (R^2^ = 0.96) for SiR-actin and -2.9° (R^2^ = 0.91) for Alexa488-phalloidin (Supplemental Fig. 3b-c). Because actin filaments are helical, bound fluorophores have cylindrical symmetry and must give θ_d_ values of either 90° or 0°. Our results agree with measured θ_d_ values of 90° for Alexa488-phalloidin ^36,37^ and with a θ_d_ value of 0° for SiR-actin, as further validated independently (Supplemental Fig. 3d). These results with actin filament fluorescence polarization validated the use of EA-TIRFM to determine the orientation of LFA-1 in migrating T cells.

**Figure 4.**
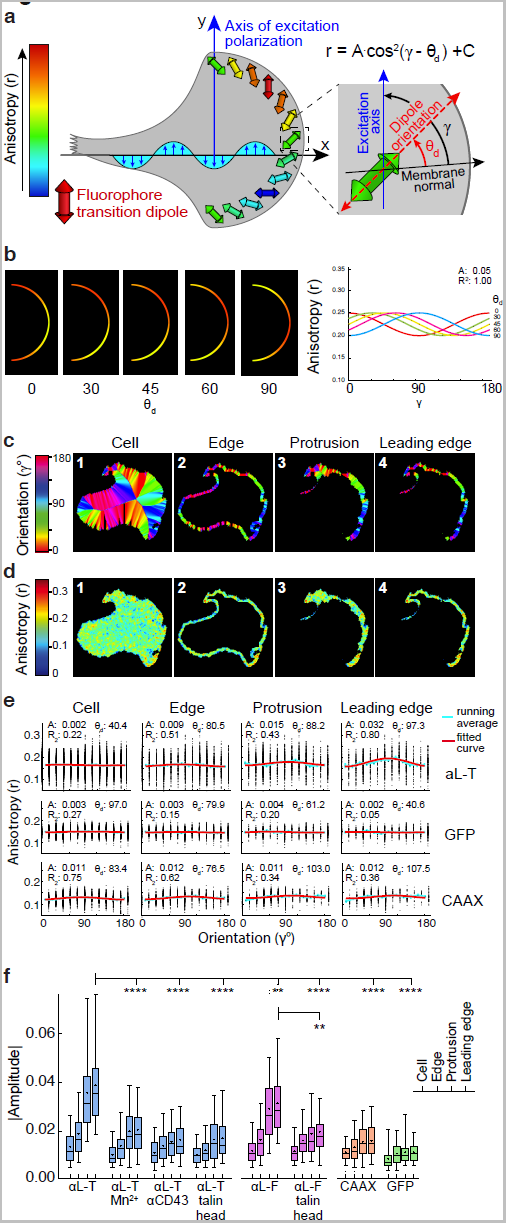
Orientation of LFA-1 in the leading edge of migrating cells by EA-TIRFM. **a.** Schematic showing relation between excitation polarization, transition dipole orientation, and emission anisotropy in EA-TIRFM. The cos^2^ function describes the relationship between orientation and anisotropy. C is isotropic background; A is absolute amplitude and reports the degree of GFP dipole alignment (which results in angular dependence); the angle between the membrane normal and the excitation axis (γ) is defined in the counterclockwise direction, as is the angle between the transition dipole and the membrane normal (θ_d_), equivalent to a phase shift in the cosine function. **b.** Simulation of ability to find a predefined dipole orientation angle relative to membrane normal in an idealized leading edge. Test data were generated with a fixed amplitude (0.05) and variable θ_d_ (0-90). The fitted curves are shown to the right and the simulated images are displayed on the left with indicated simulation θ_d_. **c-d.** Representative segmentation for a migrating Jurkat T cell expressing αL-T-GFP with corresponding maps of orientation relative to the cell midline (c) and anisotropy, *r*, (d) in the segmented regions (1-4). Scale bar = 5 μm. **e.** Emission anisotropy of segmented regions for αL-T, cytosolic GFP, and GFP-CAAX (Supplemental Figs. 4 and 7) fit to the cos2 function. Scatter plots show anisotropy vs orientation relative to the cell midline for each pixel in individual, representative cells (from c and d for αLT), the running average (blue) and fit (red). Values for the fit parameters (A = absolute amplitude, θ_d_ = phase shift, R2 = goodness-of-fit) are displayed in each plot. **f.** Absolute amplitudes of emission anisotropy fits to the cos2 relationship. Box plots show the full range (whiskers) of observations with median as line and 5-95 percentile range boxed. Kruskal-Wallis test with multiple comparison correction to αL-T leading edge data (N=206) gave p-values from left to right: <0.0001 (N=85); <0.0001 (N=52); <0.0001 (N=58); 0.0033 (N=185); <0.0001 (N=71); <0.0001 (N=83); <0.0001 (N=55) with N leading edges. See Supplemental Table 2 for more details. A two-tailed Mann-Whitney test of αL-F in absence and presence of talin head gave a p-value of 0.0059. ** p < 0.01, **** p < 0.0001.

We next measured the angular cos^2^ dependence of LFA-1-GFP anisotropy using EA-TIRFM to test the hypothesis that integrin engagement to an immobilized ligand and the cytoskeleton would cause the integrin and its associated GFP transition dipole to adopt a specific orientation relative to the direction of actin retrograde flow. Having already established that flow is normal to the leading edge of migrating cells, we tested the angular cos^2^ dependence of fluorescence anisotropy on the orientation of the leading edge relative to polarized excitation (Fig. 4a). We validated our analysis pipeline by generating ideal images of leading edge protrusions with defined dipole orientations and were able to find the correct angles (Fig. 4b). Migrating cells in movies were segmented into whole cell, edge, protruding, and leading edge (lamellipodium) regions (Fig. 4c-d, steps 1-4). Geometric shape-based orientation was determined by the angle from the cell edge to mid-region (Fig. 4c, see Methods), making it possible to test for angular dependence of anisotropy within each cell. For the most constrained construct, αL-T, angular dependence of emission anisotropy with respect to the leading edge fit the cos^2^ function (Fig. 4a). The amplitude (A), i.e. the extent of angular dependence, increased towards the leading edge (Fig. 4e, see also Supplemental Fig 4-7, Supplemental Table 2 and Movie 1). The phase shift of the maximum anisotropy of the integrin-GFP chimera relative to the angle between the membrane normal and the excitation axis in segmented leading edges, θ_d_, was 98.5° ± 37.6° for αL-T and 75.7° ± 46.6° (mean ± sd) for αL-F. Compared to the integrin-GFP fusions, fits to the cos^2^ function for anisotropy of cytosolic GFP and membrane-bound GFP (CAAX) were poor with low R^2^ values and gave low amplitudes (Fig. 4e-f, Movie 2). Activating LFA-1 independently of intracellular signaling with Mn^2+^, disrupting actin binding to the cytoplasmic domain of LFA-1 with overexpression of talin head domain, or disruption of LFA-1 engagement with the substrate on the non-integrin substrate anti-CD43 all lowered amplitude, indicative of a loss of integrin alignment (Fig. 4f). These results show that LFA-1 becomes aligned in the lamellipodium and oriented relative to the adjacent leading edge, and that the extent of alignment (A) is dependent on proper LFA-1 activation, ICAM-1 engagement, and talin linkage to the actin cytoskeleton.

We verified and extended our measurements to higher precision using a different type of fluorescence microscopy. With the Instantaneous FluoPolScope^37^, fluorophore dipoles are excited isotropically with circularly polarized laser TIRF excitation, and the emission is split four ways using polarization beam splitters and projected simultaneously onto four quadrants of a single CCD detector (Fig. 5a). Measurement of emission at four different angles (0, 45, 90, and 135°) enables determination of dipole orientation and polarization factor *p* (analogous to anisotropy *r*) in each pixel with FluoPolScope. As with EA-TIRFM, we validated the FluoPolScope with measurements on actin filaments (Supplemental Fig. 3), as recently described with the same scope^37^. We measured orientation and polarization factor of the GFP-LFA-1 emission dipole ensemble in each pixel in migrating T cells. Values in pixels were weighted by intensity in individual segments to weight all GFP-LFA-1 molecules in the segment similarly. Orientation and polarization values from multiple leading edge and cell body segments were then combined (Fig. 5b-d). Emission polarization for αL-T in leading edges was significantly higher than for αL-F in leading edges and also higher than for αL-T in the cell body or for GFP in solution or in the cytoplasm (Fig. 5c). Leading edge emission polarization factor on ICAM-1 substrates was significantly decreased by extracellular Mn^2+^ and on anti-CD43 substrates (Fig. 5c). These polarization factor results were in excellent agreement with the anisotropy results obtained by EA-TIRFM.

**Figure 5.**
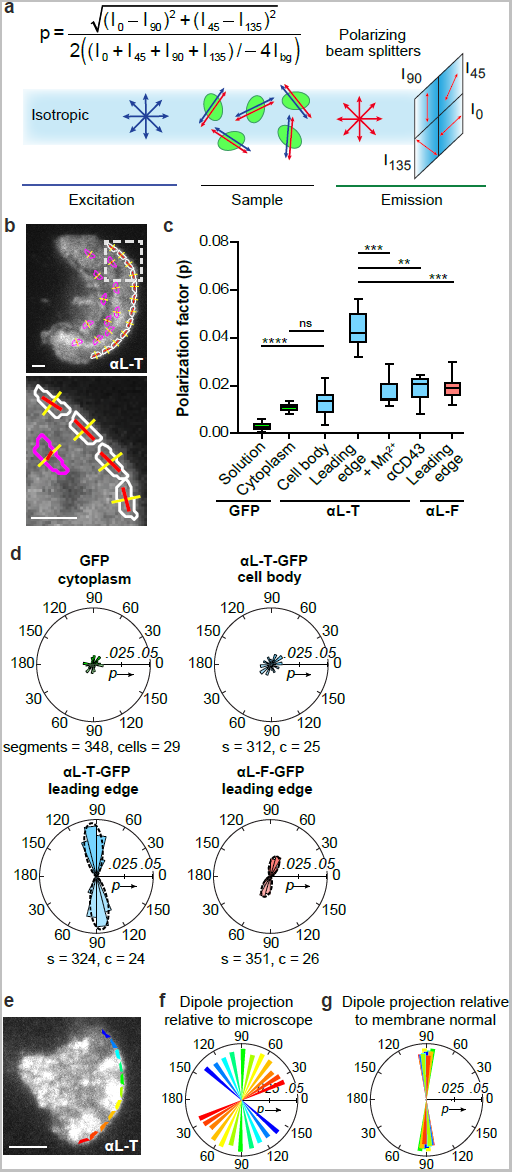
Integrin orientation along the axis of actin retrograde flow in migrating T cells measured by instantaneous FluoPolScope. **a.** Schematic of Instantaneous FluoPolScope and equation for polarization factor *p* where Ibg is background intensity. **b.** Representative total fluorescence intensity image of αL-T Jurkat T cell migrating on ICAM-1 (10 μg/ml) with overlay of segments (white = leading edge, magenta = cell body), normal to leading edge tangent (yellow), and average GFP emission dipole orientation with length proportional to polarization factor (red). Scale bars = 1μm. Panel below is enlarged from dashed area. **c.** Polarization factor of T cells migrating in random directions. Box plots show the full range (whiskers) of observations with median as line and 25-75 percentile range boxed. Kruskal-Wallis test with multiple comparison correction of leading edges gave the indicated p-values to αL-T using N number of cells shown in panel d. ** p < 0.01, *** p < 0.001, **** p < 0.0001. **d.** Radial histograms of GFP transition dipole projection in image plane relative to the membrane normal (θ_d_). Each radial histogram wedge shows mean intensity-weighted polarization factor (*p*) in 15° bins (solid outlines) and is reflected to represent the dyad symmetry axis of the dipole. Dashed propeller-shaped outlines in the lower two panels show the fit of the data to a circular Gaussian. **e-g.** e. A representative αL-T Jurkat cell migrating on ICAM-1 was segmented as shown in (b). Segments are color coded in blue-red rainbow around the leading edge and are shown in same color in f and g. Each segment is represented as a reflected wedge at the angle of its experimentally measured emission dipole projection in the microscope xy plane (f) or with respect to the membrane normal (θ_d_) (g).

Individual migrating cells show that despite the continuous change in the membrane normal from one end of the curved leading edge to the other, the emission dipole remains nearly perpendicular to the membrane normal, showing it is oriented relative to actin flow (Fig. 5b,d). When absolute dipole orientation in segments (Fig. 5e,f) is plotted relative to the membrane normal, a narrow distribution of orientations is found (Fig. 5g). We quantitated emission dipole orientation of αL-T and αL-F relative to the membrane normal (θ_d_) over 264-351 segments from 21-31 cells. Orientation was 95.4° ± 10.1° (mean±s.d.) for αL-T and 71.7° ± 17.9° for αL-F (Fig. 5d). These values are within 4° of those from EA-TIRFM, but have much smaller s.d., likely because phase shift calculations in EA-TIRFM were dependent on sampling a wide range of leading edge orientations and showed more cell to cell variation. Circular Gaussians also fit the FluoPolScope data well (propeller-shaped outlines in Fig. 5d). Taken together, the EA-TIRFM and Instantaneous FluoPolScope results show that in lamellipodia the transition dipole of the GFP moiety of GFP-LFA-1 fusions, and hence also the LFA moiety of these fusions, are oriented relative to the leading edge by retrograde actin flow.

### Translation of dipole orientation to integrin orientation on the cell surface

To translate GFP transition dipole orientation to the orientation of LFA-1 in the leading edge of migrating cells we utilized the orientation of the GFP excitation dipole relative to the GFP crystal structure measured by two independent techniques^23,38^. Based on the high anisotropy of GFP crystals, and the angular dependence of their polarized excitation and emission maxima, the excitation and emission dipoles of GFP are within a few degrees of one another^23,24^. The orientation of the GFP transition dipole was thus expressed in the same coordinate system as the GFP-LFA-1 fusions in Rosetta ensembles.

To determine the mean orientation of LFA-1 engaged to ICAM-1 and the actin cytoskeleton in migrating cells we utilized measurements of dipole orientation, and not absolute values of anisotropy or polarization factor which can be effected by how uniformly the integrins mediating cell migration are aligned to one another. It is unknown if some unaligned, unengaged integrins are present in the same regions, and whether dynamic variation in GFP orientation relative to the integrin is present, as expected from variation in orientation found in Rosetta ensembles. Thus, we relied more on αL-T-GFP than αL-F-GFP measurements, as Rosetta predicts a narrower range of dipole orientations and higher polarization factor for αL-T-GFP (Fig. 1e and Supplemental Fig. 8–10), in agreement with its higher value of experimentally measured polarization factor (Fig. 5c). Furthermore, the experimental error in αL-T-GFP dipole orientation was smaller.

Defining molecular orientation on the cell surface required constructing a frame of reference that places integrin atomic coordinates in microscope coordinates (Fig. 6a). The xy plane was defined as parallel to the microscope TIRF field, where x represents the direction of lamellipodial protrusion. The xz plane was defined by three atoms with conserved positions among integrin heterodimer ligand complexes and conformational states (Fig. 6a).

**Figure 6.**
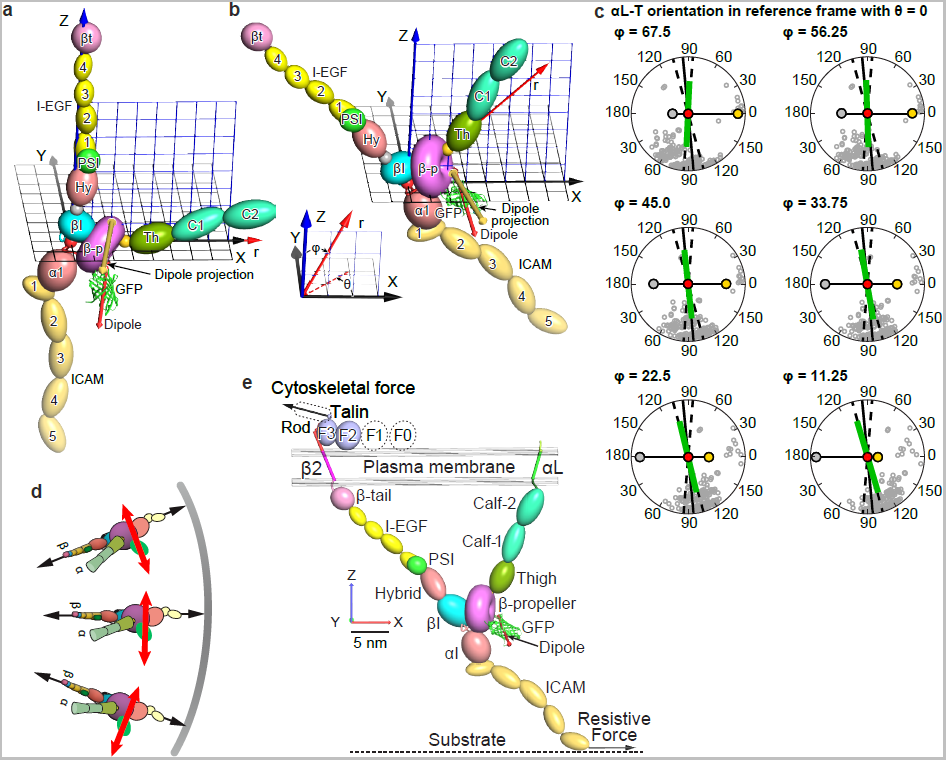
Molecular model of LFA-1 orientation on the cell surface. **a-b.** Integrin-microscope reference frame. xy and xz planes are black and blue grids, respectively. Ca atoms used to define the origin (red), x axis (gold), and xz plane (silver) are shown as large spheres. The GFP transition dipole (red) and its projection on the image plane (yellow-orange) are shown as cylinders with cones at each end. A spherical coordinate radial marker r (red arrow) is used to compare integrin orientations between the reference state with θ = 0° and ϕ = 90° (a) and integrin orientation with θ = 0° and ϕ = 45° that fits data well (b). **c.** The image plane with dipole positions of Rosetta ensemble members (lowest 40% in energy) projected from a spherical surface and shown as open grey circles. Projections are all in the reference frame with θ = 0° and with tilts in ϕ from 67.5 to 11.25°. Silver, red, and gold circles show the same integrin reference atoms as in a & b. The calculated ensemble transition dipole ^42^ is projected as a green line with length proportional to *p*. The transition dipole orientation determined by FluoPolScope is shown as black line with ± 1 s.d. shown as dashed lines. **d.** Schematic showing integrin and GFP dipole orientation relative to tensile force between the actin cytoskeleton and ICAM (arrows) in migrating cells. **e.** Model of cytoskeletal force acting on an integrin drawn to scale tilted at ϕ = 45°. Structures^11,15-17,43-46^ were assembled and rotated at domain-domain junctions known to be flexible and depicted with PyMol.

Both integrin head rotation θ in the xy plane and tilt φ relative to the z axis will affect the measured projection of the GFP transition dipole in the xy plane (Fig. 6b). The transition dipole orientation we measure is compatible with an integrin orientation of either θ = 0° or 180°; however, evidence that the cytoskeleton applies the force to the integrin β-subunit forces us to choose β retrograde of α, which is fulfilled only by θ = 0°. The experimentally determined value of θ_d_ = 95.4 ± 10.1° for αL-T is in perfect agreement with θ = 0°, as predicted by alignment by retrograde actin flow (Fig. 6c). Exploring a range of tilts (variation in φ) (Supplemental Fig. 10) shows that the transition dipole calculated from Rosetta ensemble members falls within one standard deviation of the experimentally determined θ_d_ value for φ values between 67.5° and 22.5° (Fig. 6c). The orientation predicted by αL-F-GFP is similar, but less well defined (Supplemental Fig 9). Our results define the orientation of engaged LFA-1 on cell surfaces with respect to the reference frame as within about 10° of θ = 0 and within about 25° of φ = 45.

Our results suggest that on the surface of a migrating T cell, cytoskeletal force causes the LFA-1 head to align to the direction of actin flow (Fig. 6d), and to tilt relative to the membrane normal (Fig. 6e). The integrin β-leg and ICAM-1 have flexible inter-domain junctions and the αI domain is flexibly linked to the βI domain and β-propeller domain as shown by variation among structures^2,16,17^. In contrast, orientation between the βI and β-propeller domains is uniform and enables the GFP fusions used here to report on both βI and β-propeller domain orientation. Force balance requires that tensile force straightens the flexible domain-domain linkages in this force-bearing chain like links in a tow-chain, and aligns them in the direction of force (Fig. 6e). Thus our measurements not only reveal the orientation of the integrin head near the center of this chain, but also suggest that the entire chain has an orientation similar to the path that force takes through the βI domain in the integrin head (Fig. 6e).

## Discussion

We introduce a novel method for measuring molecular orientation on cell surfaces. Little has been known about cell surface receptors built from multiple tandem extracellular domains linked to single-span transmembrane domains with respect to their orientation in the plasma membrane. Furthermore, it has been difficult to relate conformational states of isolated receptor glycoproteins to their conformation, function, and orientation on cell surfaces. We have demonstrated that integrin LFA-1 becomes aligned at the leading edge of migrating T cells and that integrin orientation correlates with actin flow orientation. Comparison of retrograde flow velocity on ICAM-1, CD43 antibody, and mixed ICAM-1/CD43 substrates demonstrated that LFA-1 slows actin retrograde flow in migrating T cells. The simplest interpretation of this result is that actin retrograde flow exerts a force on LFA-1. Measurement of force within the cytoplasmic domain of LFA-1 has directly demonstrated that force exertion requires an intact cytoplasmic domain binding site for talin, which is known to couple to the actin cytoskeleton^6^. Moreover, artificial extracellular activation of LFA-1 by Mn^2+^, which over rides regulation by actin, disrupts alignment of LFA-1. These findings are in agreement with our measurements of fluorescent dipole orientation of GFP-LFA-1 fusions, which show that the orientation of the integrin near the leading edge of migrating cells resembles that predicted by application of force by the actin cytoskeleton to the integrin β-subunit.

Our results form an important bridge between studies of forces associated with integrin adhesion and migration and structural studies on integrins. Single molecule forces measured on integrin ligands and several actin cytoskeletal adaptors have emphasized the importance of integrins in transmitting force between the extracellular matrix and the cytoskeleton^4,5^. Recently, forces have also been measured within integrin cytoplasmic domains in T cells migrating on ICAM-1 substrates ^6^. Force is exerted on the cytoplasmic domain of the LFA-1 β-subunit, but not its α-subunit, and is dependent on binding to ICAM-1 on the substrate and intact binding sites in the β-subunit cytoplasmic domain for talin and kindlin. Our results here show that the forces exerted on integrins are sufficient to align them, and that the specific orientation found for integrins at the leading edge of migrating cells is consistent with the orientation predicted for force application by the cytoskeleton to the integrin β-subunit cytoplasmic domain that is transmitted through the integrin and resisted by a ligand bound to a substrate.

In contrast to force measurements in cells, crystal, EM, NMR, SAXS, and neutron scattering structures of integrins are determined in the absence of force. Elegant structures have revealed intact integrins, integrin ectodomains and their complexes with ligands, integrin transmembrane domains, and integrin cytoplasmic domains and their complexes with intracellular effectors that link or inhibit linkage to the cytoskeleton^2,3,7,8,39,40^. The demonstration here that we can use fluorescence microscopy to define a specific orientation for integrin atomic structures on the surface of migrating cells now makes it inescapable to discuss integrin structural biology in the context of force application by actin retrograde flow. For example, flexibility between many of the domains in integrins enables them to straighten and align with their domain-domain junctions parallel to the force. Flexibility in poorly structured residues that link the ectodomain to the plasma membrane enables integrin tilting^15^ as suggested by our measurements here. Moreover, the integrin β-subunit transmembrane domain tilts when separated from the α-subunit transmembrane domain, as occurs upon integrin activation^2,7,8^, consistent with the tilt suggested here.

While almost every paper on an integrin complex discusses how binding of a ligand or effector may regulate integrin activation by selecting among integrin conformational states, the implications of complex formation for tensile force transmission from the cytoskeleton that could select among integrin conformational states is discussed by few workers in the field. Perhaps one of the most important structural biology implications of our findings on integrin orientation on cell surfaces is the strong support it provides for the cytoskeletal force model of integrin activation^2,6,9–12^. Among the three integrin conformational states, only the extended-open state has high affinity for ligand^2,41^. When considering conformational equilibria between states, tensile force stabilizes equilibria by a potential energy equal to the force times the difference in extension along the force bearing axis in each state. Since the extended-open conformation is the most extended of the three states, this high affinity state is favored by tensile force. Stabilizing high integrin affinity for extracellular ligand by force application by the actin cytoskeleton provides a simple and elegant mechanism for coordinating cytoskeletal activity inside the cell with binding to ligands in the extracellular environment during cell migration^6,11^. The mechanisms that regulate actin cytoskeleton dynamics are highly complex, and the coordination by tensile force transmission of ligand binding with actin cytoskeletal motion not only avoids the need to have a regulatory mechanism for integrin activation that duplicates actin regulatory mechanisms but also enables efficient coordination between the actin cytoskeleton and integrins to provide cellular traction precisely where cytoskeletal force is exerted. Thus, our results place molecular understanding of integrin structure and function within the context of integrin function in cell migration and in linking the extracellular environment to the actin cytoskeleton. Since 22 of the 24 mammalian integrin αβ heterodimers have β-subunits that link to the actin cytoskeleton, the findings here with integrin LFA-1 are expected to be of wide relevance among integrins.

## Author contributions

This project was initiated at the Woods Hole Physiology Course. PN, TIM, SM, CMW and TAS designed the research. TT, SM, CMW, RO, and TAS supervised the project. PN and TAS designed and made the GFP-integrin constructs. TIM performed imaging experiments and analyzed data. PN wrote image processing and analysis code and analyzed data. SBM and JMK set up polarization microscopes and RO advised on analysis. NK, TAS, and DB performed Rosetta modeling. TAS constructed the integrin-microscope reference frame and defined the GFP transition dipole orientation within Rosetta models. TAS and SBM analyzed Rosetta data. TIM, TJL, and JCW performed SIM experiments and analysis. SBM, JMK and VS contributed to image data analysis. PN, TIM, and TAS drafted the manuscript. All authors discussed the results and commented on the manuscript.

## Acknowledgements

We thank Nikon Instruments and Andor Technology for use of imaging equipment, Thomas Holder at Schrodinger for Pymol/Python scripts and advice to create cgo representations of ellipsoids, torus, and dipoles, and Einat Schnur, Gabriel Billings, Darius Vasco Köster, and Amy Gladfelter for help, advice and discussion. Supported by the Lillie Research award from Marine Biological Laboratory and the University of Chicago (CMW, TAS, SM, TT), NIH 5R13GM085967 grant to the Physiology Course at Marine Biological Laboratory, HHMI Summer Institute at Marine Biological Laboratory (SM), NIH CA31798 (TAS, PN, TIM), NIH GM100160 (TT, SM), NIH GM092802 (DB, NK), NIH GM114274 (RO, SM) National Center for Biological Sciences-Tata Institute of Fundamental Research (SM, JMK), JC Bose Fellowship and HFSP Grant RGP0027/2012 (SM), NHLBI Division of Intramural Research (CMW, VS), Swedish Research Council VR 524-2011-891 Fellowship (PN), Swedish Society for Medical Research SSMF Fellowship (PN).

## Methods

### Integrin-GFP constructs

EGFP or moxGFP^1^ were inserted into the β3-β4 loop of blade 4 of the αL integrin β-propeller domain^2^. Integrin Gly residues adjacent to GFP were mutated to Ala or Gln residues for helix propensity as indicated, linkers were added for flexibility, or GFP residues were deleted for less flexibility as follows. N and C-terminal insertion sites are shown with residues for integrin wt or mutant sequences in plain text, linkers in bold, and GFP underlined: αL-GFP-F, EPQG MVSKGEELF…MDELYK GGHW; αL-GFP-L, EPQG**SGSG** MVSKGEELF…MDELYK **GSGS** GGHW; αL1, EPQA EELF…MD AQHW; αL2, EPQA EELF…MDE AQHW; αL3, EPQA EELF…MDEL AQHW; αL4, EPQA ELF…MDELY AQHW; αL5, EPQA LF…MDELY AQHW. The αL3 construct worked best during functional testing of αL1- αL5 and was used throughout this study with the name αL-GFP-T.

Integrin α and β-subunit cDNA were made using three-segment (A,B,C) overlap PCR with wild type human ɑL cDNA and either pEGFP-N1 (Clontech) or moxGFP^1^ (for ɑL-GFP-T) as sources for GFP cDNA. After the three segments had been made and stitched together through PCR (Accuprime Pfx, high-fidelity polymerase, ThermoFisher), the complete A–C sequence and the wild type αL-pcDNA3.1 plasmid were cut with restriction enzymes (New England Biolabs) and ligated together with T4 ligase (Roche) after dephosphorylation (rAPID alkaline phosphatase, Roche) and purification (Qiagen) of the linearized plasmid. The overall plasmid integrities were verified with size matching of multi-site single restriction enzyme digestion compared to virtual digest patterns (Serial Cloner) and the inserts were verified by full sequencing. Surface expression of the αL-GFP constructs was validated by transient co-expression with β2 in 293T cells. For ɑL-GFP-F the primers used were: A1: 5´-AGA TGT GGT TCT AGA GCC ACC ATG AAG GAT TCC TGC-3´; A2: 5´-TGA ACA GCT CCT CGC CCT TGC TCA CCA TGC CCT GTG GCT CTT GGA AC-3´; B1: 5'- AGT GCT GCT GTT CCA AGA GCC ACA GGG CAT GGT GAG CAA GGG CGA G -3'; B2: 5'- ATG GAT TGT CTG GAC CTG GCT CCA GTG TCC TCC CTT GTA CAG CTC GTC CAT GCC -3'; C1: 5'- ATC ACT CTC GGC ATG GAC GAG CTG TAC AAG GGA GGA CAC TGG AGC CAG -3'; C2: 5'- ACT CTT AGT AGC GGC CGC TCA GTC CTT GCC ACC ACC -3'. Primers for ɑL-GFP-L were: A1: 5'- AGA TGT GGT TCT AGA GCC ACC ATG AAG GAT TCC TGC -3'; A2: 5'- AGC TCC TCG CCC TTG CTC ACC ATG CCA GAT CCA GAG CCC TGT GGC TCT TGG AAC -3'; B1: 5'- AGT GCT GCT GTT CCA AGA GCC ACA GGG CTC TGG ATC TGG CAT GGT GAG CAA GGG CGA G -3'; B2: 5'- TGT CTG GAC CTG GCT CCA GTG TCC TCC GCT GCC TGA GCC CTT GTA CAG CTC GTC CAT GCC -3'; C1: 5'- ATC ACT CTC GGC ATG GAC GAG CTG TAC AAG GGC TCA GGC AGC GGA GGA CAC TGG AGC CAG -3'; C2: 5'- ACT CTT AGT AGC GGC CGC TCA GTC CTT GCC ACC ACC -3'. Primers for ɑL-GFP-T were: A1: 5'- AGA TGT GGT TCT AGA GCC ACC ATG AAG GAT TCC TGC -3'; A2: 5'- CAC CAG AAT AGG GAC CAC TCC AGT AAA CAG TTC CTC AGC CTG TGG CTC TTG GAA CAG CAG - 3'; B1: 5'- GGC CGA GTG CTG CTG TTC CAA GAG CCA CAG GCT GAG GAA CTG TTT ACT GGA GTG GTC CC -3'; B2: 5'- CCA TGG ATT GTC TGG ACC TGG CTC CAG TGT TGA GCA TAC AGC TCA TCC ATT CCG TGG GTG -3'; C1: 5'- GCT GCT GGA ATC ACC CAC GGA ATG GAT GAG CTG TAT GCT CAA CAC TGG AGC CAG GTC CAG -3'; C2: 5'- ACT CTT AGT AGC GGC CGC TCA GTC CTT GCC ACC ACC -3'. See Extended Data Table 1 for details on amino acid sequence and constructs used for simulations. Other constructs used were: Lifeact ^3^ fused with mCherry or mNeonGreen, talin head fused to mApple, and mApple with CAAX sequence added.

### Reagents

Wild-type soluble ICAM-1-His6 (D1-D5) was expressed in 293 cells and purified on Ni-NTA agarose^4^. Human SDF1-α was from R&D System. Cytochalasin D was from Santa Cruz. Blebbistatin was from AbCam. Phalloidin-Alexa 488 was from Invitrogen. Anti-CD43 was from ebiosciences. Glass-bottom dishes and plates were from Mattek. Leibovitz’s L-15 medium and RPMI-1640 medium were from Life Technologies. The reagents for the lentiviral Gateway system were from Life Technologies. Nucleofector Kit V was from Lonza.

### Cells

Jurkat T cells (clone E6.1) were cultured in RPM1-1640 medium with 10% FBS in 5% CO_2_ and supplemented with 3 μg/ml puromycin and/or 1 μg/ml blasticidin if they had been lentivirally transduced.

### Lentiviral transduction of cells

The Gateway system from Invitrogen was used to create lentiviral constructs. The integrin constructs were inserted either into pLX302 or pLX304. Virus was produced in 293T cells by co-transfecting the lentiviral plasmids with psPAX2 and CMV-VSV-G. Virus in supernatants was concentrated using Lenti-X. Jurkat cells were transduced and selected using puromycin (pLX302) or blasticidin (pLX304).

### Live imaging

Glass-bottom dishes or plates were adsorbed overnight at 4° C with 10-20 μg/ml ICAM-1 in carbonate buffer (pH 9.6), followed by blocking at 37° C with 1% BSA in L-15 medium for 30-60 min, and washing with base imaging media consisting of L-15 supplemented with 2 mg/ml glucose. Cells were suspended in base medium supplemented with 100 ng/ml SDF1-α. Before imaging, cells were added to the dish or well on the microscope held at 37 °C and allowed to settle.

### Fixed cell imaging

Cells were prepared as for live imaging and allowed to migrate at 37 °C for 30 min. Inhibitors were added for inhibitor-specific times prior to fixation: DMSO, 1:2000, 30 min; cytochalasin D, 100 nM, 15 min; blebbistatin, 100 uM, 30 min. Fixation with an equal volume of paraformaldehyde at a final concentration of 2% was for 10 min at 37 °C. After washing with PBS, cells were imaged as for live samples.

### Actin purification, polymerization, labeling and glass fixation

Actin was purified from chicken breast following the protocol from Spudich et al^5^. The monomeric form was maintained in G-buffer (2 mM Tris Base, 0.2 mM ATP, 0.5 mM TCEP-HCl, 0.04% NaN_3_, 0.1 mM CaCl_2_, pH 7.0) on ice. For actin polymerization, the G-actin was mixed with G-buffer and 10% v/v of 10x ME buffer (100 mM MgCl_2_, 20 mM EGTA, pH 7.2) to obtain an actin concentration of 10 μM and incubated for 2 min to replace G-actin bound Ca^2+^ ions with Mg^2+^. Next an equal amount of polymerization buffer was added to induce F-actin polymerization at a final actin concentration of 5 μM in KMEH (50 mM KCl, 2 mM MgCl_2_, 1 mM EGTA, 20 mM HEPES, pH 7.2) supplemented with 2 mM ATP and 1 mg/ml BSA. After 20-30 min incubation, the F-actin was labeled by addition of 500 nM Phalloidin-Alexa488 (Invitrogen) and/ or 1 μM SiR (Cytoskeleton Inc) and incubation of 10 min at room temperature. Then, F-actin was sheared by pipetting up and down 10 times, diluted in KMEH and transferred to a 0.01% Poly-L-Lysine coated glass bottom dish at final concentration of 10 nM. After 15 minutes of incubation, unbound actin filaments were washed away with KMEH buffer and the samples were imaged in a 100X 1.49 NA TIRF microscope. For Poly-L-Lysine coating, 0.01% PLL was aseptically coated onto the surface of No 1 glass coverslips and rocked gently to ensure even coating. After 5 minutes, the excess solution was removed by aspiration and the surface was rinsed with tissue culture grade water and left drying under laminar flow for at least 2 hours before use.

### Emission anisotropy total internal reflection fluorescence microscopy (EA-TIRFM)

EA-TIRFM images were acquired using the TIRF mode on a Nikon Eclipse TiE inverted microscope equipped with a motorized TIRF illuminator (Nikon, USA) and a motorized stage (TI-S-ER motorized stage with encoders; Nikon, USA) fed by a multi-wavelength (405 nm [15-25mW], 488 nm [45-55mW], 561 nm [45-55 mW], 640 nm [35-45 mW]) polarization-maintaining fiber coupled monolithic laser combiner (Model MLC400, Agilent Technologies). This arrangement generates a polarized TIRF evanescent field at the sample plane^6^.

Images were collected with a fixed magnification using a 100X Plan Apo 1.49 NA TIRF objective (Nikon, USA) fitted with a Perfect Focus System (PFS3; Nikon, USA) and a 1.5X tube lens to yield a final pixel size corresponding to 109 nm. The typical TIRF illumination depth using 488 nm was 150-200 nm. Band-pass emission filters (ET525/50, ET600/50 and ET700/75; Chroma Technology Corp, USA) were mounted onto a motorized turret below the dichroic mirror (405/488/561/638 TIRF Quad cube; Chroma Technology Corp, USA).

Emission from the polarized evanescent TIRF field was split into constituent *p* and *s*-polarized components using a high performance nano-wire grid polarizing beam splitter (TR-EMFS-F03; Moxtek Inc.,USA). The resulting parallel and perpendicular components were imaged with separate, orthogonally placed iXon Ultra 897 EMCCD cameras (Andor Technology, Belfast, Northern Ireland) using the TuCam two-camera imaging adapter (C-Mount Version [S-CMT]; 1X Magnification [TR-DCIS-100]; Andor Technology, Belfast, Northern Ireland). Images were acquired using the Nikon Imaging Software (NIS Elements Advanced Research; Nikon, USA) with a dual-camera plug-in using the electron multiplying (EM) gain mode.

### Instantaneous FluoPolScope

A custom microscope using opto-mechanics from Newport Corp was built on an optical table. Laser beams (Coherent Sapphire 488 nm, 20mW and Melles-Griot 561 nm, 25mW) were routed through custom optics and focused on the back focal plane of a 100x 1.49 NA objective (Nikon 100x ApoTIRF 1.49NA). The objective was placed on a Piezo Z-collar (PIP-721 PIFOC) for precise focusing. Laser beams were circularly polarized using a combination of a half wave plate and quarter wave plate (Meadowlark Optics). To achieve isotropic excitation within the focal plane and along the optical axis of the microscope, the circularly polarized laser beam was rapidly rotated (300-400Hz) in the back focal plane of the objective with a large enough radius to achieve total internal reflection at the specimen plane. Dual-band dichroic mirror (Semrock Di01-R488/561) was used to separate laser lines (reflected) and emissions corresponding to GFP and mCherry (transmitted). The specific emission channel was selected using bandpass filters mounted in a filter wheel (Finger Lakes instruments). A quadrant imaging system as described in Mehta et al^7^ was used for instantaneous analysis of fluorescence emission along four polarization orientations at 45° increments (I0, I45, I90, I135). Dual-channel imaging of live cells was performed using Micro-Manager (version 1.4.15). All images were acquired using an EMCCD camera (Cascade II: 1024; Photometrics, Tuscon, AZ) operated in the 5MHz readout mode with EM gain.

### Structured illumination microscopy (SIM)

3D-SIM data were collected on a DeltaVision OMX V4 Blaze system (GE Healthcare) equipped with a 60x / 1.42 N.A. plan Apo oil immersion objective lens (Olympus), a 488 nm diode laser, and an Edge 5.5 sCMOS camera (PCO). Image stacks of ~2-3 μm (fixed) or ≤1 μm (live) were acquired with a z-step of 125 nm and with 15 raw images per plane (five phases, three angles). Spherical aberration was minimized using immersion oil matching^8^.

### Image processing and analysis

#### EA-TIRFM

Image processing and analysis was mainly carried out using MATLAB 2014a. Functions handling all steps of the image processing were developed into a semi-automatic software package with optional manual steps for image registration. Stepwise, images were imported from the original files and sorted into channels; all metadata were extracted and saved; image registration was carried out with one of three options: manual reference image of submicron beads initialization, automatic reference image initialization or automatic registration using cell images; G factors were calculated daily based on fluorescein solution images; images were G factor corrected; and background was masked by thresholding at a value 3 standard deviations above background, where the background intensity distribution is estimated by fitting the “left half” of a Gaussian function (the portion below its mean) to the left shoulder of the image intensity histogram. This mask was then used to find and subtract the average background intensity on a frame-by-frame basis for each channel. For all anisotropy calculations the data was pre-filtered with a 3x3 intensity-weighted average applied to all pixels. To minimize artifacts from division of small integers, only pixels with intensities above 4 times the background standard deviation of the current frame were used; finally, anisotropy was visualized using a heat map. Intensity-weighted anisotropy (r) of each group of nine pixels was calculated and displayed in the central pixel through the relationship:

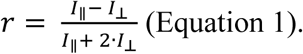

In this equation the difference between parallel and perpendicular intensity is divided by total intensity where perpendicular intensity is added twice to account for the two planes perpendicular to the parallel plane in three-dimensional fluorescence emission.

For image analysis the initial step was to identify cells. Briefly, since multiple cells could be present in the same image and have different fluorescent expression, the background masks described above was used to initialize cell segmentation independently for each potential cell area. These regions were slightly expanded (5-pixel dilation operation) to make sure that background was included to maintain contrast for increased robustness of the algorithm. Either active contour segmentation (energy minimizing that separates between foreground and background) or intensity distribution-based threshold segmentation (similar to the background masking described above) was used to produce an initial cell mask. Mathematical morphology (a closure operation with a radius of 1 pixel, small object removal, and filling of holes) was applied to further refine these masks, producing accurate cell outlines. For live imaging data, a four-dimensional bounding box (3D space plus time dimension of at least 5 frames) was used to make sure that cells were consistently segmented during the movie. For edge segmentation, the cell mask was eroded by 10 pixels, and then inversely combined with the original mask to generate an edge mask. For protrusion detection, the difference between the cell masks in neighboring frames were evaluated with a four-dimensional bounding box (positive area of 2000 pixels and at least 5 frames) and were stored as individual protrusion masks. Leading edge segmentation was carried out by combining edge and protrusion masks for positive protrusions. Signal-to-noise ratio (SNR) was determined for each segmented region,

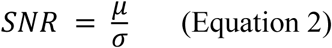

where μ is the mean intensity of the perpendicular channel and σ is the standard deviation of the background intensity in that channel. Any region with a SNR lower than 5 was excluded from analysis (Extended Data Fig. 1c). Given the variable cell shapes, especially between frames and in protrusive regions, an orientation mapping algorithm was devised that would assign relative orientation values in a reproducible manner. It is based on calculating the vector away from the edge for each pixel in cell masks. For circular cells, it yields an orientation axis of 0 to 180°, that falls along the polarization axis with clockwise assignment of orientation values (Extended Data Fig. 7d). For irregular shapes such as polarized cells, the orientation assignment is not always comparable across cells, only within. This also means that for non-circular objects the orientation values are not absolutely correlated with the polarization axis and introduces variation in phase shift estimates across cells, but with consistent angular dependency estimates (amplitude). The cell masks were smoothed with a 3-pixel radius closure operation followed by an Euclidian distance transform of the inverse mask,

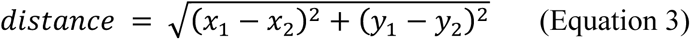

with x and y pixel coordinates. A numerical 2-dimensional gradient is calculated from the distance transform,

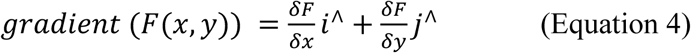

and the inverse tangent is used to return the relative orientation value for each pixel. To assess angular dependence, the orientation values were binned to the closest 10 or 15 degrees and then the orientation and anisotropy values for each pixel were fitted to a cos^2^ function,

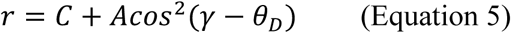

and the absolute amplitude of this is used as a measure to report the degree of angular dependence. The angle between the membrane normal and the excitation axis (γ), is defined in the counterclockwise direction, as is the angle between the transition dipole and the membrane normal (θ_d_), equivalent to a phase shift in the cosine function. To verify whether it is possible to obtain a correct θ_d_ from assigning orientations to an anisotropy map, and fitting these to Equation 5, simulated images with known variables were used (Fig. 4b). In all cases, the fits were perfect (R^2^=1.00), and the correct amplitude and phase shifts were found.

To get an alternative measure of angular dependence, a Fourier-based analysis (Fast Fourier Transform) was used on the running average of orientation relative to anisotropy (Extended Data Fig. 4–7). The DC term was removed by subtracting the mean anisotropy value and the amplitude value from the second bin (or first frequency, equivalent to a basic wave) was recorded.

#### Instantaneous FluoPolScope

Image analysis was performed using custom code developed in MATLAB 2014a. Algorithms are available upon request. The four polarization resolved quadrants of the integrin-GFP channel were cropped, registered, and throughput-normalized as described in ^7^. The total intensity image of each cell was segmented along the leading edge into 1000 nm × 500 nm masks with the long axis tangential to the membrane. The cell body was segmented using masks with the same long axis angle relative to the microscope frame of reference, to ensure no effect of segmentation orientation, and distributed around the cell body more than 1000 nm from the leading edge. Segments were digitized, and membrane normal (yellow line in Fig. 5b) computed as perpendicular to the segment long axis. Background polarization and excitation imbalance were determined for each cell from an approximately 0.28 μm square (4×4 pixels) within the center of each cell. Background-corrected polarization-resolved intensities (I0, I45, I90, I135) were then summed over each segment. These sum intensities per segment were used to compute dipole orientation (*θ*) and polarization factor (*p*) per segment as follows:

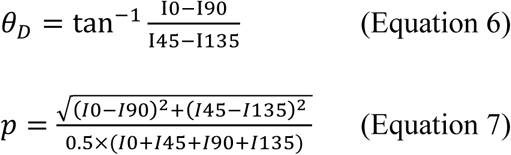

For each segment, GFP dipole orientation relative to the membrane normal, θ_d_, was calculated as the angle in the counter-clockwise direction from the membrane normal.

#### Structured illumination microscopy

Super-resolution images were computationally reconstructed from the raw datasets with a channel-specific measured optical transfer function (OTF) and a Wiener filter constant of 0.002-0.003 using softWoRx 6.1.3 (GE Healthcare). Live-cell datasets were collected at 37˚C using objective and stage-top heaters (GE Healthcare) and fixed datasets were collected at room temperature. Live cell image acquisition was optimized to keep peak laser intensity below ~30 W/cm^2^ and exposure times below 10 ms (≤ 2 sec per Z-stack) to minimize motion-induced reconstruction artifacts.

Actin flow velocity and direction was determined using the Flow-J optical flow analysis^9^ plugin within the Fiji image processing package^10,11^. The Lucas and Kanade algorithm^12^ was applied to each live structured illumination time series to calculate the velocity vector for each pixel in each frame. The velocity vectors were sampled along the leading edge for each substrate condition to determine actin flow speed and angle relative to the microscope frame of reference X axis. Concurrently, the angle of the membrane tangent of each segment was measured. The angles were subtracted to determine the angle of actin flow relative to the cell membrane. Mean actin velocity was calculated and conditions compared for statistical difference using the Mann-Whitney test.

Kymographic analysis to determine actin flow velocity was performed as described in Comrie et al^13^. SIM movies of T cells expressing Lifeact-mNeonGreen were analyzed using the Fiji image processing package to generate a vertical kymograph traversing the cell leading edge and lamellipodium. The flow rate was calculated based on the slope of deflection of F-actin from the vertical direction^13^ (Supplemental Fig. 2).

### Estimation of GFP dipole orientation relative to integrin

Low energy orientations between the inserted GFP and integrin were efficiently sampled using Rosetta. Rosetta found low energy conformations for “loop” sequences at the two integrin-GFP junctions. The remainder of integrin was rigid and GFP was rigid except for solvent-exposed sidechains. Conformations of the junction loops were found that permitted loop closure and prevented rigid body clashes followed by further loop relaxation and sidechain optimization to minimize energy^14^. Integrin sequences within two residues of the insertion site, linker residues, and GFP residues that vary in position or are disordered in GFP structures (residues 1-5 (MVSKG) and 228-238 (GITLGMDELYK) were included in the loop regions that were subjected to backbone optimization. For computational efficiency, only β-propeller residues 330-483 were included in the α_L_β_2_ model; other domains of the α-subunit and the β-subunit were too distal to clash with GFP, as confirmed when full length models were built subsequent to modeling the GFP- α_L_β_2_ fusions. The αI domain can vary markedly in orientation relative to the β-propeller domain in which it is inserted; therefore, we chose the most physiologic orientation available, from an α_X_β_2_ ectodomain crystal structure in which the internal ligand of the αI domain binds to a pocket at the β-propeller interface with the βI domain, concomitantly with activation of the high affinity, open conformation of the αI domain^15^. A complete LFA-1 ectodomain model was built by superimposing the β-propeller domain from an α_L_β_2_ headpiece crystal structure^16^ and the αI domain from a complex of its high affinity state with ICAM-1^17^ onto the cocked α_x_β_2_ crystal structure^15^.

For each integrin-GFP fusion construct, Rosetta output an ensemble of structures that effectively sampled low energy GFP-integrin orientations. Longer loop junctions typically enabled a larger number of loop closures enabling larger ensembles; however, no closures were found for αL5 suggesting its loops were too short. The ensemble of GFP orientations was visualized by superimposition on the integrin in the integrin-microscope reference frame. Dipole orientations were calculated in spherical coordinates. Models were ranked according to total energy and compared for angular distribution. The two lower energy quintiles had a more restricted range of dipole orientations; therefore we used the 40% lowest energy models as the ensemble for calculations of ensemble dipole orientation and polarization factor.

The first chemically plausible orientation (within the planar ring system of the amino acid residues that fuse to form the fluorophore) for the GFP excitation dipole was determined from the polarized light absorption spectra by GFP crystals and had to account for the four distinct GFP molecules present with different dipole orientations in the crystal lattice and four mathematically possible solutions to the equations used to define excitation dipole orientation^18^. Subsequently, the equations were corrected and the solution refined^19^. Additionally, it was realized that previously reported visible pump /IR probe measurements could be used to calculate the orientation of the fluorophore transition dipole relative to carbonyl stretch vibrational transition moments that are well defined spectroscopically and assigned to carbon-oxygen bonds between atoms well characterized for their positions in GFP crystal structures^19^. The transition dipole orientations calculated by these two independent methods were in good agreement, and authors of the latter publication (X. Shi and S.G. Boxer) kindly provided the transition dipole orientation for GFP chain B in the coordinate system of the high-resolution structure PDB ID code 1w7s as a line with slope x = -0.026, y = 0.871, and z = 0.439. A line with this slope drawn through the hydroxyl oxygen atom of the chromophore closely matched a line with α = 6.5° in Fig. 6 of Shi et al^19^ and was approximated in integrin-GFP ensembles as a line drawn through the Val-112 N atom and the average of the positions of the Asn-146 C and Ser-147 O atoms in the GFP moiety. Rosetta ensemble transition dipole orientations and polarization factors were calculated as described^20^.

### Integrin orientation on the cell surface

In the extended-open conformation in which integrins bind ligands, the interface of the α-subunit β-propeller and β-subunit βI domains, which form the head and bind external ligands (as with α_V_β_3_) or internal ligands (as with α_L_β_2_), faces away from the integrin legs which connect to the transmembrane and cytoplasmic domains. To orient the liganded integrin in this manner, we first skeletonized the ligand-bound integrin into three key Cα atom points. The ligand-point is at the strongly bound Asp of RGD or internal ligand Glu of αI integrins. Two junction points are near sites of pivoting movements between the head and legs, yet are sufficiently inward in the head to show little variation in position among independent integrin-ligand complexes or among conformational states, and are sufficiently conserved in position to enable comparison among distinct integrin heterodimers. Furthermore, these points move little in steered molecular dynamics simulations (SMD)^21^. The α-junction point is at the C-terminus of the α-subunit β-propeller domain (Arg-438 in α_V_, Gln-451 in α_IIb_, Arg-588 in α_L_, or Arg-597 in α_X_) that connects to the flexible thigh domain. The β-junction point is at the N-terminus of the βI domain (Pro-111 in β_3_ or Pro-104 in β_2_) that connects to the hybrid domain. These junction points are between β-strands in adjacent domains, each of which are central strands in their β-sheets and are thus highly force-resistant.

Cartesian and spherical coordinate reference frames were defined to enable dipole orientations measured in microscopes, the orientation of the transition dipole in GFP, and the orientation of GFP with respect to the integrin in Rosetta ensembles to be used to define integrin orientation on cell surfaces. The coordinate system is defined by the three Cα atoms in the integrin and ligand described in the preceding paragraph. Because force passes through the junction between the ligand and integrin, and force balance requires that tensile force cause them to pivot toward alignment with the direction of force exertion, the ligand-point is used to define the origin. The line between the ligand point and the α-junction point defines the X axis. The β-junction point defines the XZ plane and lies near the Z axis. These positions in turn define the XY plane, which lies parallel to the microscope image plane and the plasma membrane of the cell adhering through integrins to ligands on the substrate.

As the reference frame is constructed with the ligand and head junction points in the XZ plane and optimally orients the integrin head toward the substrate, the integrin will remain close to this plane when it is tilted by cytoskeletal force, as shown by SMD^21^. Spherical coordinates are useful for defining the orientation of the integrin and GFP transition dipole relative to the Cartesian reference frame. X, Y, Z positions in Cartesian coordinates are defined in spherical coordinates with the radial distance r and angles θ and φ. Integrin and dipole orientations are defined relative to the X axis with the α-junction point and r lying on this axis such that θ=0° and φ= 90°. The orientation of r is defined by its angle φ with the Z axis and its angle θ with the X axis when projected on the XY plane. Similarly, the orientation of the GFP transition dipole when projected on the microscope imaging plane in our fluorescence microscopy experiments is measured as θ_d_ relative to the direction of lamellipodial movement in the direction θ = 0° (normal to the leading edge). The reference frame is such that when actin retrograde flow is in the direction θ = 180°, the traction force model for integrin activation predicts that 1) the line between the ligand and the α-junction points tilts toward the Z axis with a decrease in φ and 2) the projection of this line on the XY plane has θ = 0°, as in the reference frame.

---

**Supplemental Figure 1.**
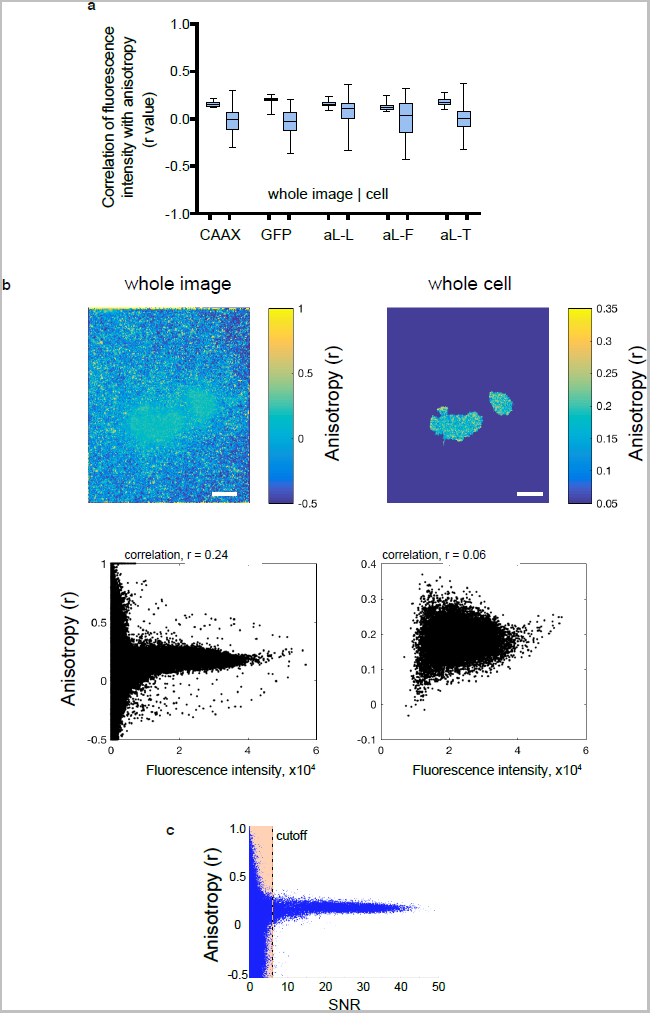
Correlation of anisotropy with fluorescence intensity or signal to noise ratio (SNR). **a.** Correlation r values between total fluorescence intensity (combined polarization channels, 2 perpendicular + 1 parallel) and anisotropy. Correlation values are shown for whole images (left box) and segmented cells from images (right box). Box plots show the full range (whiskers) of the data and 25-75% range (boxes) with median as line. From left, number of images N = 33, 55, 206, 185, and 36. **b.** Representative image showing anisotropy (upper) and plot of total fluorescence intensity versus anisotropy (lower), with whole image data (left) and segmented cell data from the same image (right). Note difference in anisotropy scales. Scale bars are 5 μm. **c.** Representative plot of anisotropy vs SNR in a non-segmented image. At low SNR, the anisotropy values are uncertain and therefore an SNR cutoff threshold (SNR>5) was used for all EA-TIRFM analysis.

**Supplemental Figure 2.**
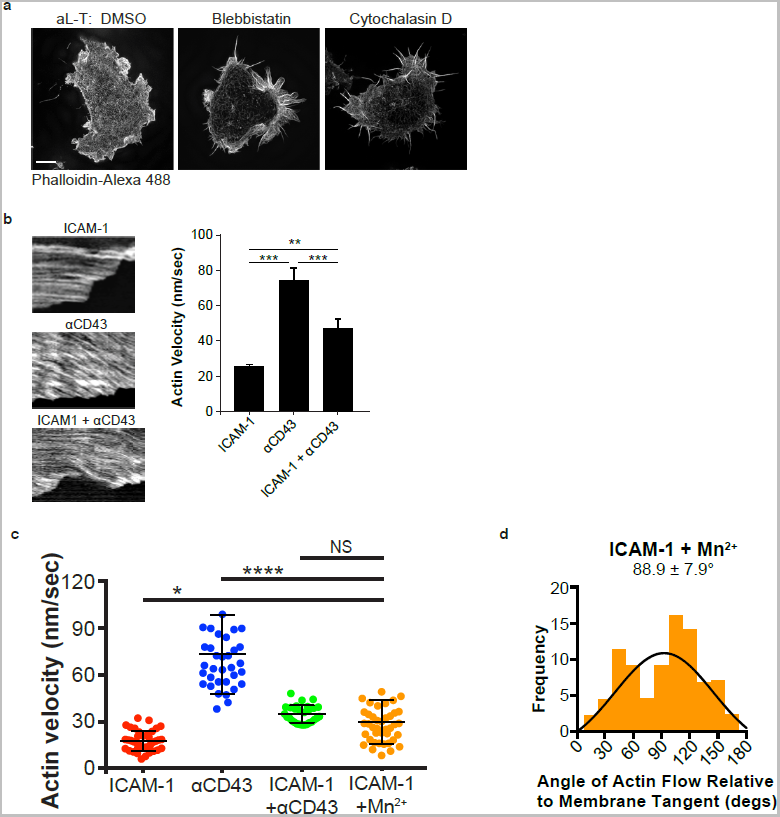
Additional SIM actin data. **a.** Representative super-resolution SIM images of the actin cytoskeleton in fixed Jurkat T cells migrating on ICAM-1 as labeled with Alexa 488-phalloidin treated with DMSO, 100 μM blebbistatin or 100 nm cytochalasin D. Scale bar = 2 μm. **b.** Kymograph analysis of retrograde actin flow and representative kymographs (see Methods). ICAM-1, N=30; aCD43, N=24; ICAM-1+αCD43, N=27. Bars show mean ± SEM. Two-tailed Mann-Whitney tests; p < 0.01 (**), p < 0.0001 (****). **c.** Leading edge actin flow velocity from optical flow analysis. Plots show full range of the data. Bars show mean ± SD. Two-tailed Mann-Whitney tests; p < 0.5 (*), p < 0.0001 (****). ICAM-1, N=39; αCD43 N=38; and ICAM-1+ anti-CD43, N=25; ICAM-1+Mn_2+_, N=41. Some data shown in Fig. 3 is repeated here for comparison. **d.** Actin flow direction relative to tangent of leading edge membrane. Bins of 15° are shown with gaussian fit and mean ± SEM. ICAM-1, N=39; aCD43, N=37; ICAM-1+αCD43, N=26; ICAM-1+Mn2+, N=42.

**Supplemental Figure 3.**
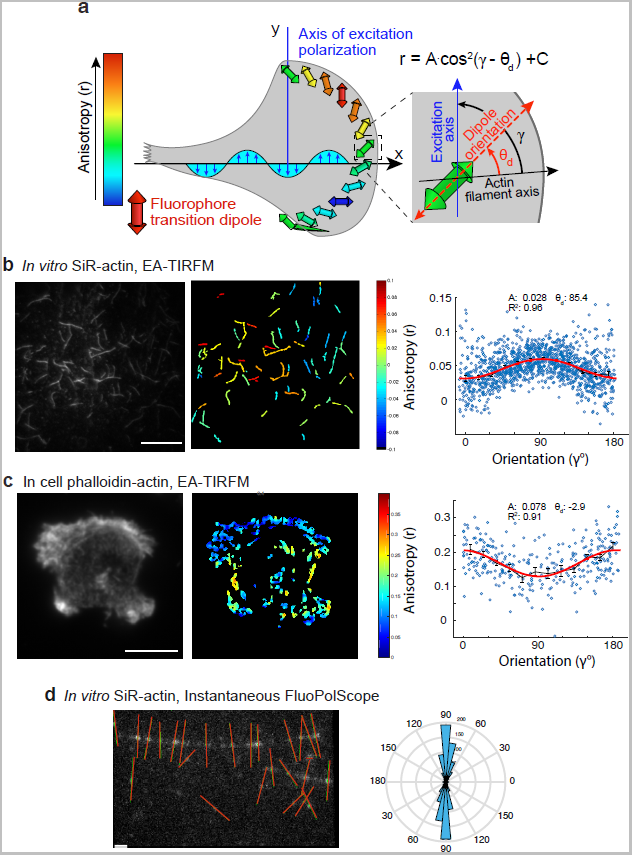
Orientation of actin filaments determined with EA-TIRFM and Instantaneous PolScope. **a.** Schematic showing relation between excitation polarization, transition dipole orientation, and emission anisotropy in EA-TIRFM. Repeated from Fig. 4a. **b.** Left, representative example of *in vitro* SiR-actin filaments with fluorescence, anisotropy and scatter plot of anisotropy vs orientation angle with data on filaments from 15 images using EA-TIRFM. Right, running average is shown as a black line and the fit curve using cosine function (see Methods and a) is shown in red. Scale bar is 10 μm. **c.** Representative example of actin phalloidin in fixed Jurkat T cells with total fluorescence (left), anisotropy (middle) and plot of anisotropy vs orientation angle (right) with data from 6 cells using EA-TIRFM, running average shown as a black line, and fitted curve using cosine function (see Methods and a) shown in red. Scale bar is 5 μm. **d.** Left, lines indicate ensemble dipole orientation of SiR-actin probe bound within different segments of actin filaments in vitro using Instantaneous FluoPolScope. Right, angular distribution of the ensemble dipole orientation relative to the filament orientation (θ_d_). Radial axis is the number of filaments which fall within each angular bin. Mean orientation ± full width at half-maximum: 91.4° +-14°. N (number of filaments analyzed) = 643. Scale bar is 1 μm.

**Supplemental Figure 4.**
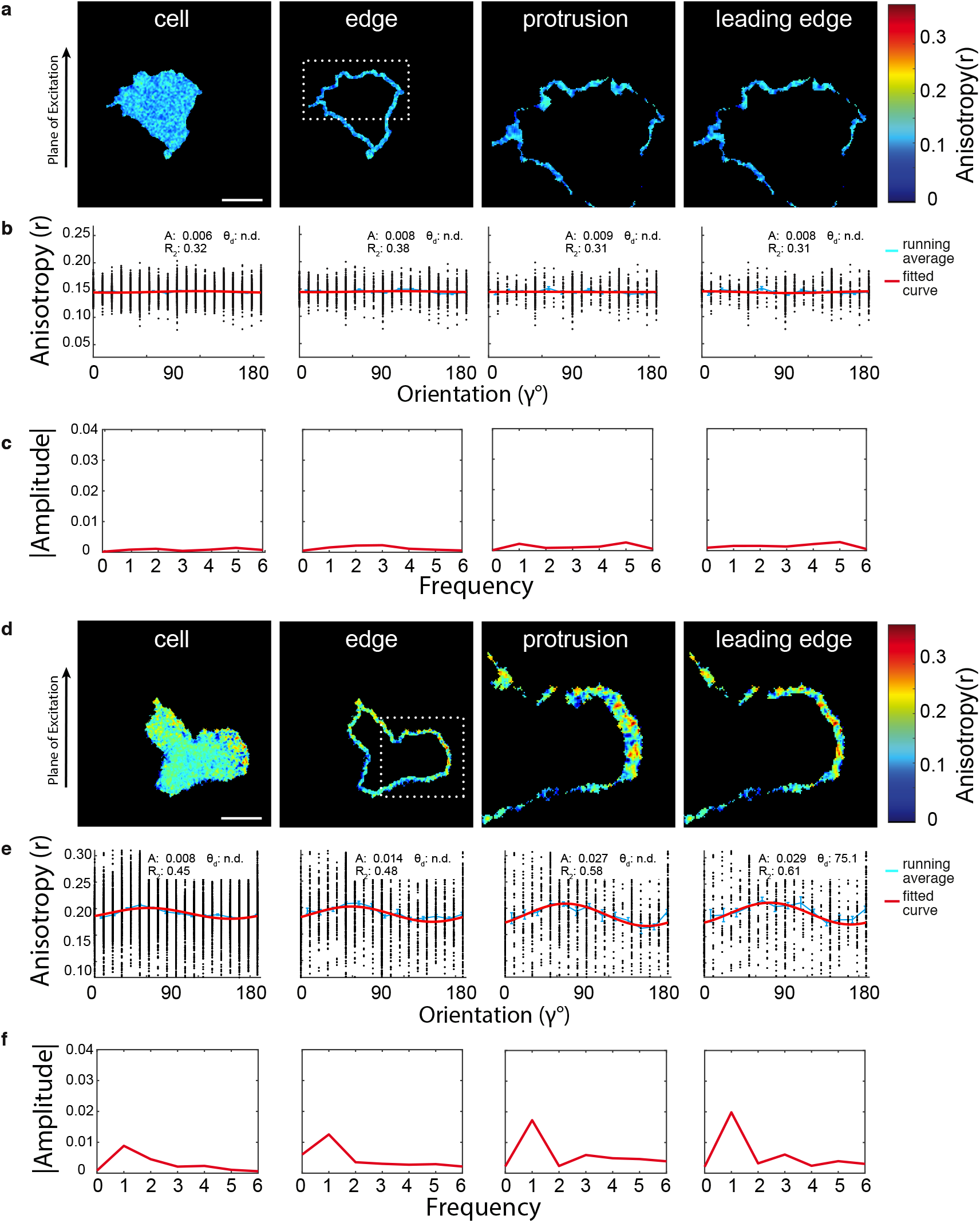
Angular dependence analysis – Cytosolic GFP (a-c) and aL-GFP-F (d-f) **a, d.** Representative example of migrating Jurkat T cell expressing cytosolic GFP (a) and aL-GFP-F (d), respectively. Each cell is segmented into whole cell, edge, protrusion and leading edge regions (see Methods). Scale bar is 5 μm. **b, e.** Scatter plot of anisotropy vs orientation angle with data from a and d. Running average is shown as a blue line and a fitted curve using cosine function (see Methods) is shown in red with mean values from fit above. See Extended Data Table 2 for tabulated values. **c, f.** Power spectrum plot from Fourier transform of data from b and e. Absolute amplitude of frequency peaks is plotted on y-axis.

**Supplemental Figure 5.**
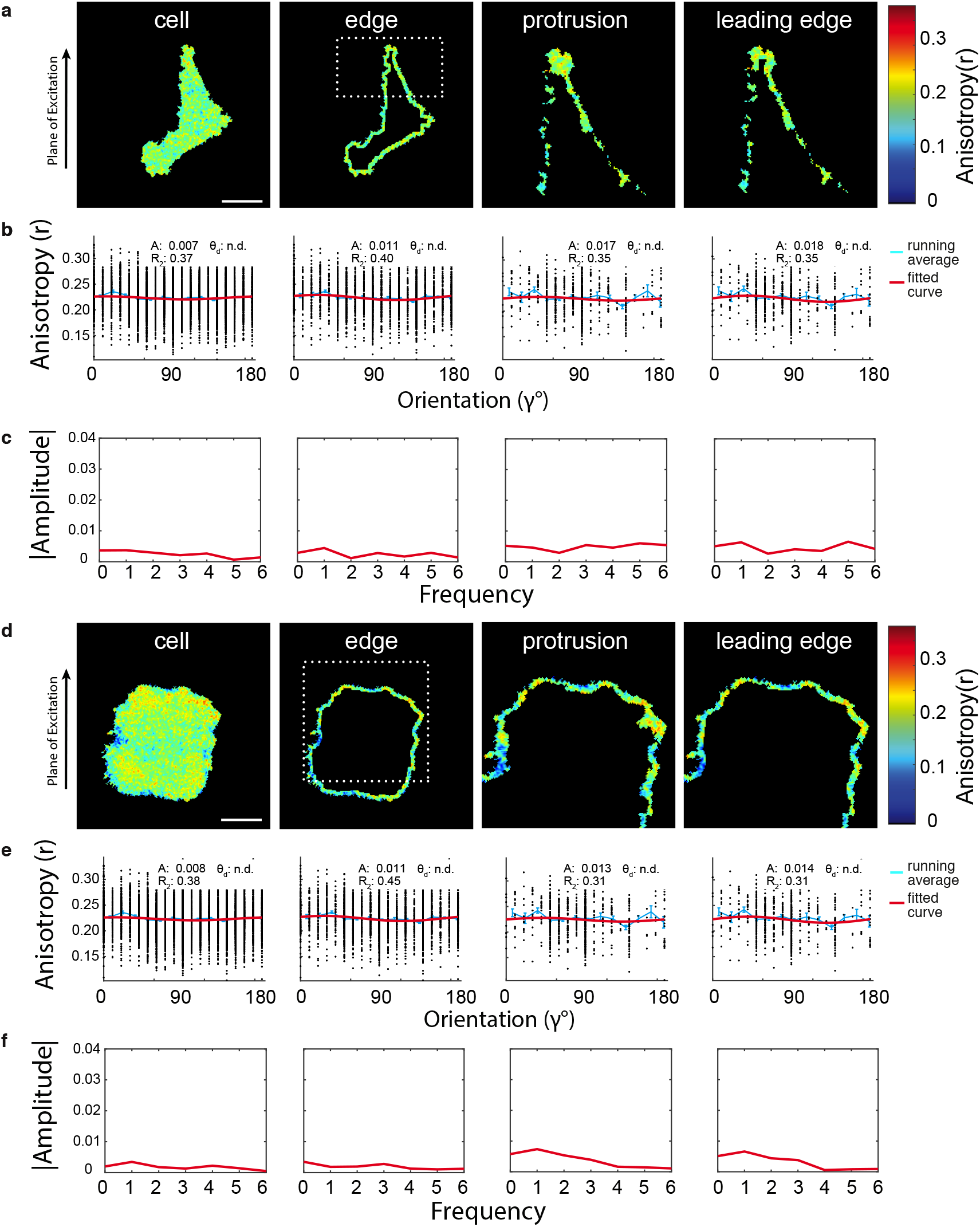
Angular dependence analysis – aL-GFP-T with Mn2+ (a-c) or on anti-CD43 (d-f) **a, d.** Representative example of migrating Jurkat T cell expressing aL-GFP-T with Mn2+ (a) or on anti-CD43 (d), respectively. Each cell is segmented into whole cell, edge, protrusion and leading edge regions (see Methods). Scale bar is 5 μm. **b, e.** Scatter plot of anisotropy vs orientation angle with data from a and d. Running average is shown as a blue line and a fitted curve using cosine function (see Methods) is shown in red with mean values from fit above. See Extended Data Table 2 for tabulated values. **c, f.** Power spectrum plot from Fourier transform of data from b and e. Absolute amplitude of frequency peaks is plotted on y-axis.

**Supplemental Figure 6.**
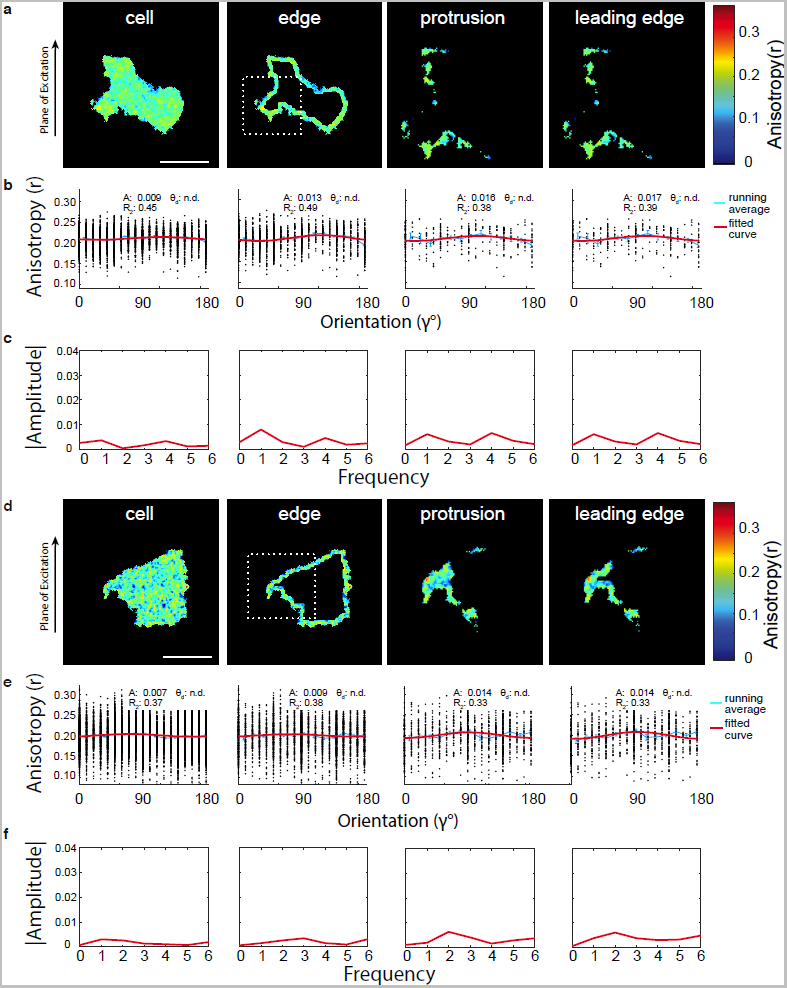
Angular dependence analysis – talin head overexpression with aL-GFP-F (a-c) and aL-GFP-T (d-f) **a, d.** Representative example of migrating Jurkat T cell overexpressing talin head with aL-F-GFP (a) or aL-GFP-T (d). Each cell is segmented into whole cell, edge, protrusion and leading edge regions (see Methods). Scale bar is 5 μm. **b, e.** Scatter plot of anisotropy vs orientation angle with data from a and d. Running average is shown as a blue line and a fitted curve using cosine function (see Methods) is shown in red with mean values from fit above. See Extended Data Table 2 for tabulated values. **c, f.** Power spectrum plot from Fourier transform of data from b and e. Absolute amplitude of frequency peaks is plotted on y-axis.

**Supplemental Figure 7.**
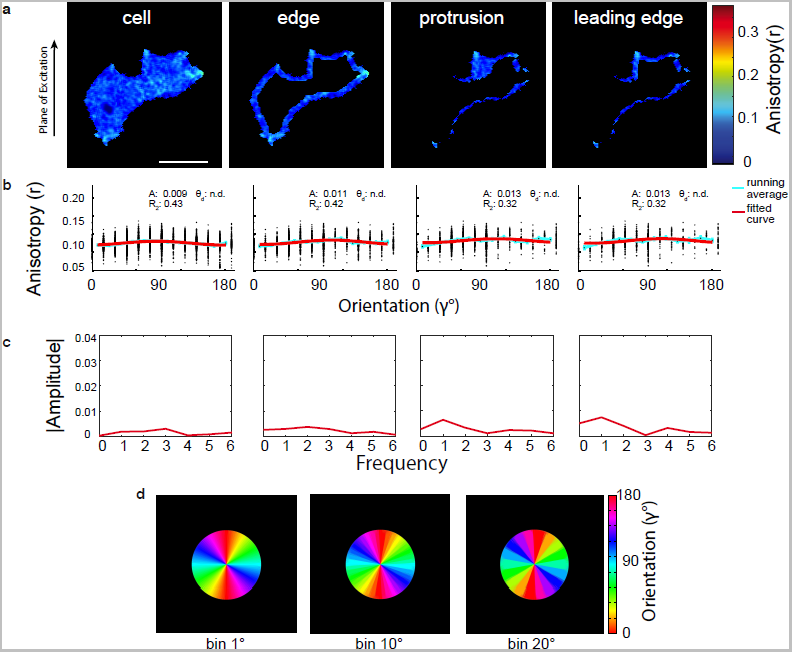
Angular dependence analysis of CAAX and orientation algorithm simulation data. **a.** Representative example of migrating Jurkat T cell expressing CAAX-GFP. The cell is segmented into whole cell, edge, protrusion and leading edge regions (see Methods). Scale bar is 5 μm. **b.** Scatter plot of anisotropy vs orientation angle with data from a and d. Running average is shown as a blue line and a fitted curve using cosine function (see Methods) is shown in red with mean values from fit above. See Extended Data Table 2 for tabulated values. **c.** Power spectrum plot from Fourier transform of data from b. Absolute amplitude of frequency peaks is plotted on y-axis. **d.** The images show three circles where relative orientation was determined with orientation values binned to 1, 10 or 20 degrees. The algorithm is described in the Methods section.

**Supplemental Figure 8.**
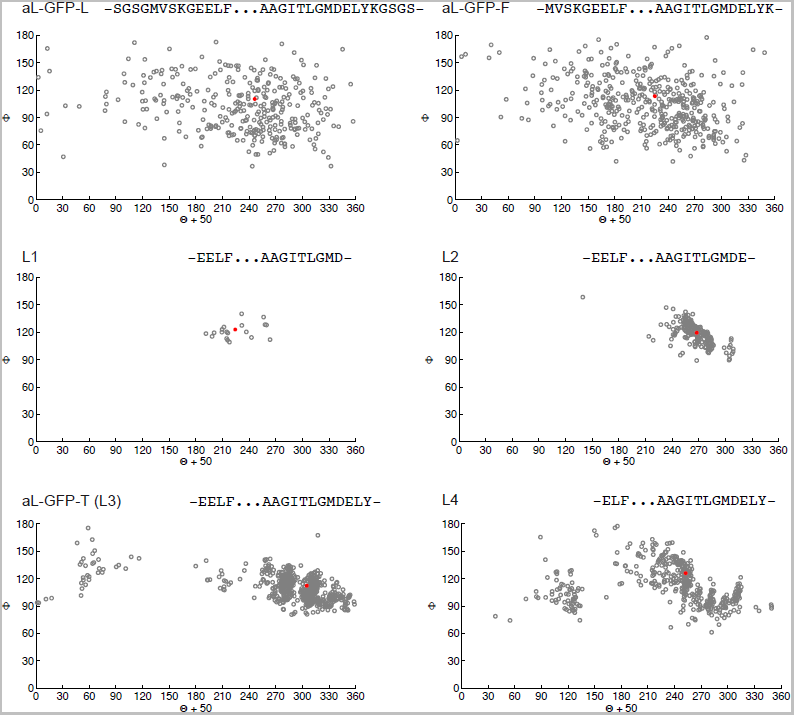
Transition dipole orientation in Rosetta ensembles: dependence on residues included at the GFP-integrin fusion junctions. Each open circles show GFP dipole orientations in the integrin-microscope frame of reference for Rosetta ensemble members. The red dot is the centroid of all ensembles. Residues at integrin juctions with N and C-terminal portions of GFP are shown above each plot, with construct names as described in Extended Data Table 1.

**Supplemental Figure 9.**
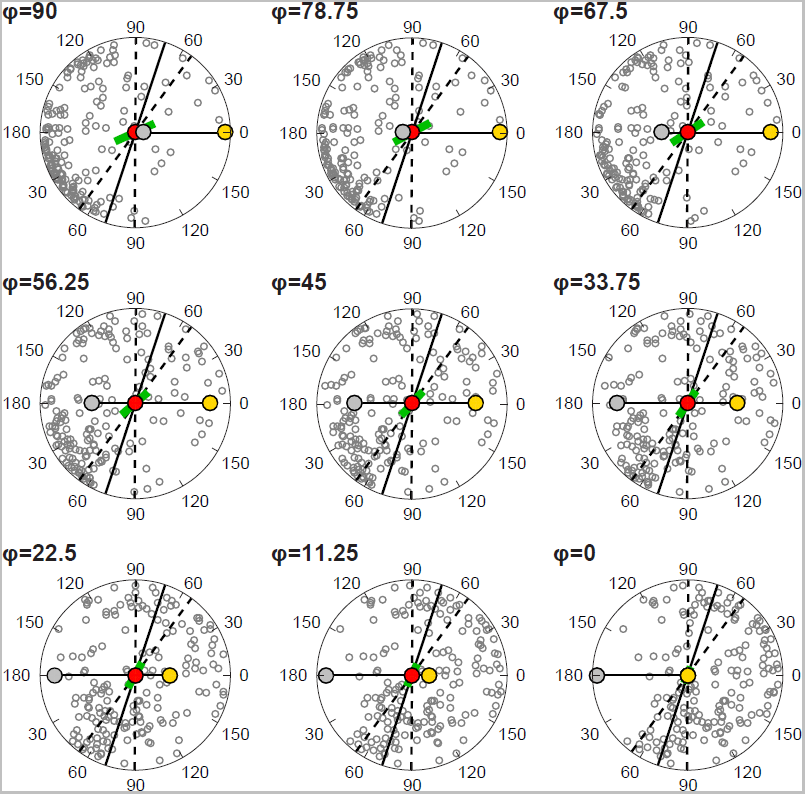
Effect of aL-F tilt on dipole orientation in the XY plane of the integrin-microscope reference frame. GFP transition dipole orientation for each Rosetta ensemble member (lowest 40% in energy) was represented by its position on the surface of a sphere in spherical coordinates in the integrin-microscope reference frame. The projection of each ensemble member in the XY plane of the reference frame at θ = 0 and φ at the indicated tilt relative to the Z axis is shown as an open circle. The calculated ensemble transition dipole is shown as a green line with orientation θ and length scaled to the polarization factor such that *p*=1 at the radius of the sphere that is projected as a circle. Red, gold, and silver circles represent the projected positions of the three integrin atoms that define the reference frame and correspond to key ligand, α-leg junction, and β-leg junction residues (see main text Fig. 4h). FluoPolScope experimental dipole projection measurements are shown with mean orientation (solid black line) ± 1 sd (dashed black lines).

**Supplemental Figure 10.**
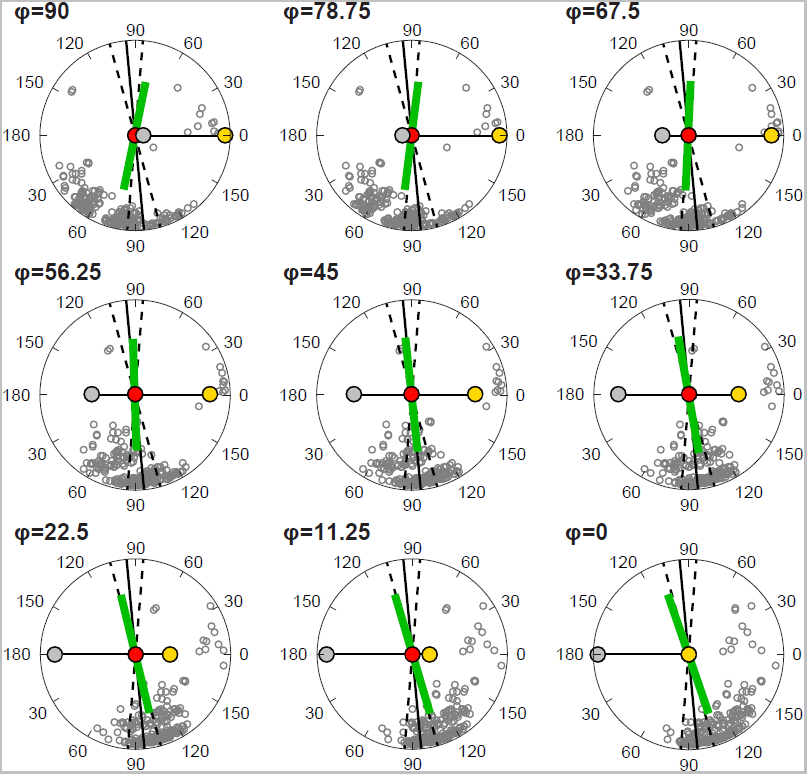
Effect of aL-T tilt on dipole orientation in the XY plane of the integrin-microscope reference frame. GFP transition dipole orientation for each Rosetta ensemble member (lowest 40% in energy) was represented by its intersection with the surface of a sphere in spherical coordinates in the integrin-microscope reference frame. The projection of each ensemble member in the XY plane of the reference frame at θ = 0 and φ at the indicated tilt relative to the Z axis is shown as an open circle. The calculated ensemble transition dipole is shown as a green line with orientation θ and length scaled to the polarization factor such that *p*=1 at the radius of the sphere that is projected as a circle. Red, gold, and silver circles represent the projected positions of the three integrin atoms that define the reference frame and correspond to key ligand, α-leg junction, and β-leg junction residues (see main text Fig. 4h). FluoPolScope experimental dipole projection measurements are shown with mean orientation (solid black line) ± 1 sd (dashed black lines). Note: each dipole was shown only in one direction from the origin to represent the asymmetry of the GFP molecule in which the dyad-symmetric dipole is present. To represent this asymmetry of GFP, the projections include θ values from 0 to 360°. However, the dyad symmetry of the dipole in the plots is represented by graphing two series of θ values from 0 to 180°, and by reflecting the calculated and experimentally observed transition dipoles.

**Supplemental Table 1.**
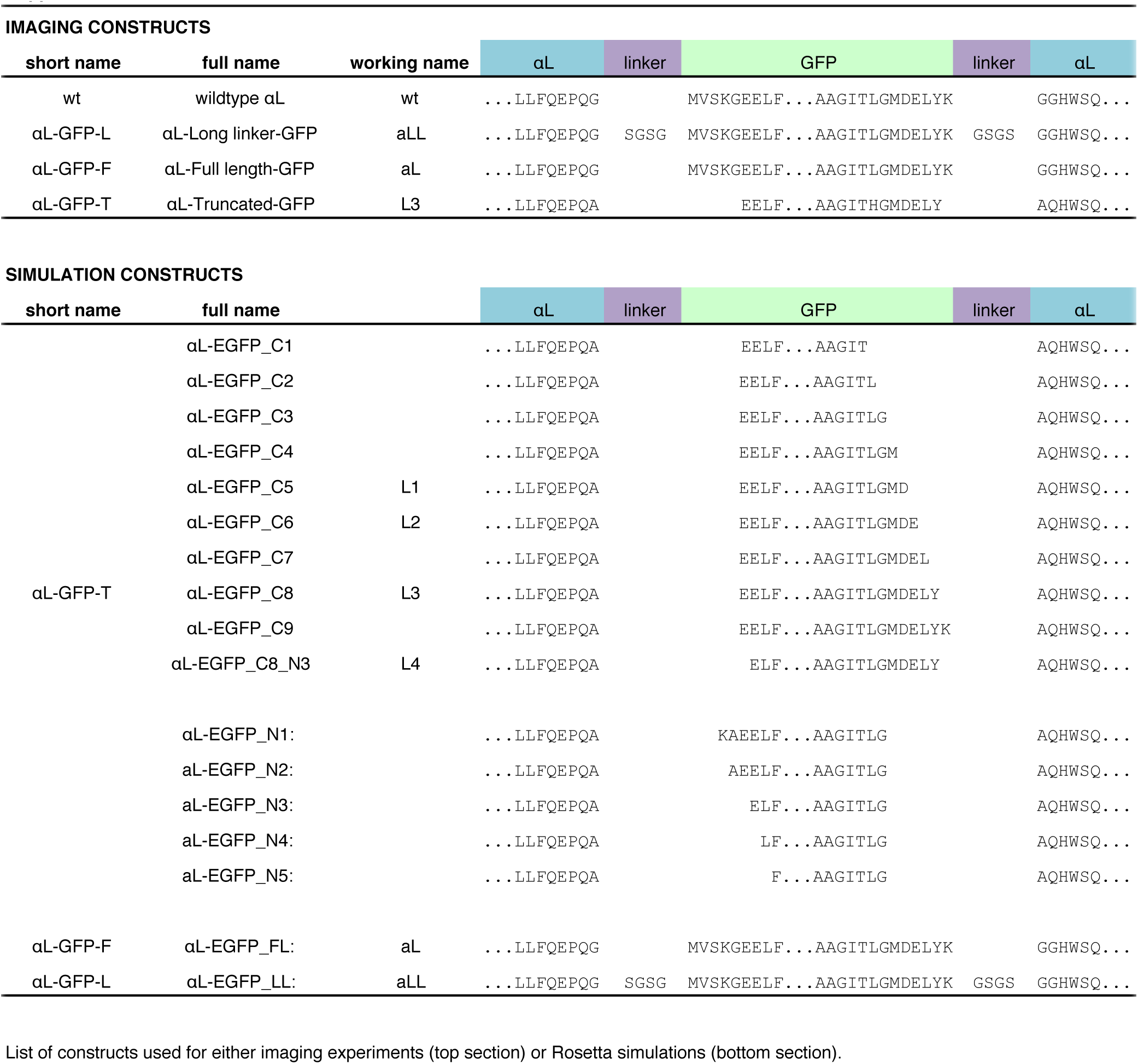
List of αL-GFP constructs.

**Table S2.**
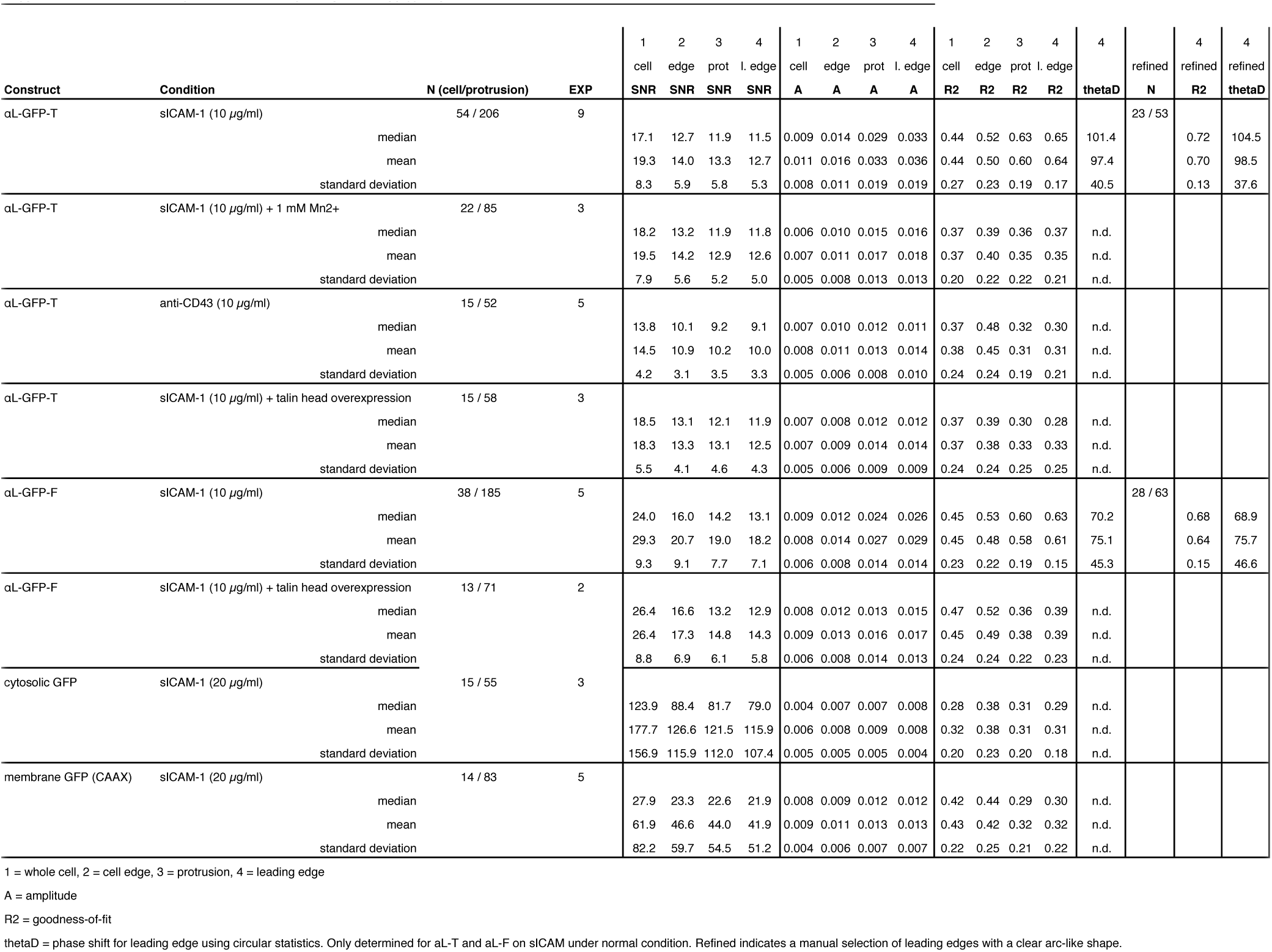
Summary of data obtained by fitting anisotropy (r) vs geometric orientation.

